# Spatial coupling between DNA replication and mismatch repair in *Caulobacter crescentus*

**DOI:** 10.1101/2020.07.13.200147

**Authors:** Tiancong Chai, Céline Terrettaz, Justine Collier

## Abstract

The DNA mismatch repair (MMR) process detects and corrects replication errors in organisms ranging from bacteria to humans. In most bacteria, it is initiated by MutS detecting mismatches and MutL nicking the mismatch-containing DNA strand. Here, we show that MMR reduces the appearance of rifampicin resistances more than a 100-fold in the *Caulobacter crescentus Alphaproteobacterium*. Using fluorescently-tagged and functional MutS and MutL proteins, live cell microscopy experiments showed that MutS is usually associated with the replisome during the whole S-phase of the *C. crescentus* cell cycle, while MutL displays an apparently more dynamic association with the replisome. Thus, MMR components appear to use a 1D-scanning mode to search for rare mismatches, although the spatial association between MutS and the replisome is dispensible under standard growth conditions. Conversely, the spatial association of MutL with the replisome appears as critical for MMR in *C. crescentus*, suggesting a model where the β-sliding clamp licences the endonuclease activity of MutL right behind the replication fork where mismatches are generated. The spatial association between MMR and replisome components may also play a role in speeding up MMR and/or in recognizing which strand needs to be repaired in a variety of *Alphaproteobacteria*.

## INTRODUCTION

It is critical for cells to replicate their genome precisely and efficiently. This process is inherently accurate due to the high fidelity of replicative DNA polymerases and their associated proofreading activities. On rare occasions, however, bases can still be mis-incorporated by the replisome, leading to potentially deleterious mutations if not repaired before the genome gets replicated again during the next cell cycle. Fortunately, nearly all cells possess a DNA mismatch repair (MMR) system that can detect and correct such errors, increasing the fidelity of DNA replication by 50-1000 folds (1). Thus, MMR prevents the appearance of drug resistances, genomic instability or cancer development in a variety of organisms, from bacteria to humans (2–5).

The MMR process was initially discovered in the *Escherichia coli Gammaproteobacterium* where it is initiated by MutS, MutL and MutH (6–9). According to the so-called “sliding clamp model”, MutS bound to ADP searches for mismatches on newly synthesized DNA. When it detects a mismatch, MutS exchanges its ADP for ATP, leading to a conformational change allowing it to diffuse along the DNA until it recruits MutL. MutS is then recycled back into its searching MutS-ADP mode through its ATPase activity. MutS-activated MutL recruits the MutH endonuclease, which can recognize which of the two DNA strands is not yet methylated by the orphan Dam DNA methyltransferase, corresponding to the newly synthesized strand that needs to be nicked and repaired (10). Then, MMR must take place within minutes after base mis-incorporation by the replisome, or else the methylation-dependent signal might be lost due to fast methylation of the newly synthesized strand. The speed of mismatch detection may be affected by the physical association between replisome and MMR components, so that MMR takes place where mismatches are created (11,12). Once the mismatch-containing DNA strand has been cut by MutH, it is unwound by the UvrD helicase and degraded by several single-stranded exonucleases. The resulting ssDNA gap is then replicated by the DNA polymerase III before being sealed by the DNA ligase (1,13,14).

While MutS and MutL homologs can be found in most organisms, it is not the case for MutH and Dam homologs that are found in only a subset of *Gammaproteobacteria*, uncovering the limits of using *E. coli* as the only model organism to study the MMR process (15). In most other prokaryotic and eukaryotic organisms, MutL carries the endonuclease activity cutting newly synthesized DNA during the MMR process (1,4,16). The *Bacillus subtilis* Gram-positive bacterium emerged as an alternative and informative model to dissect the complex interactions between MMR proteins, replication proteins and DNA in live bacterial cells using methylation-independent MMR processes (15). Single molecule microscopy observations suggested that MutS is usually dwelling at the replisome in *B. subtilis* cells, consistent with constant exchange of MutS molecules at the replisome when searching for mis-paired bases (8,17). Following mismatch detection, MutS appears to transiently diffuse away from the replisome as a sliding clamp recruiting MutL, which is then licensed to nick the nascent DNA strand by the DnaN β-clamp of the DNA polymerase (15,18). It still remains unclear how MutL recognizes the newly replicated DNA strand and whether a helicase is then needed to locally unwind the DNA before the nicked DNA strand gets degraded by the WalJ exonuclease (15). While the interaction of MutL with DnaN appears as mostly accessory in *E. coli*, it was shown to be critical for MMR in *B. subtilis* (11,18,19). Furthermore, *in vitro* assays using purified *B. subtilis* MutL and DnaN demonstrated that the endonuclease activity of MutL is dependent on its interaction with the β-sliding clamp (19). Whether the importance of dynamic spatial associations between replisome and MMR components are a general feature of MutH-independent MMR processes in bacteria or just a specific mechanism of action found in firmicutes or Gram-positive bacteria, remains an open question (15). Thus, there is a need to characterize MMR systems in more diverse bacterial species to address this important issue.

The *Caulobacter crescentus Alphaproteobacterium* appears as an interesting model to study MutH-independent MMR beyond Gram-positive bacteria, since the replication of its chromosome has been the subject of intensive studies over the last decades (20). Unlike most bacteria, its cell cycle is easily synchronizable, it shows clear G1/S/G2-like phases and replication never re-initiates within the same cell cycle, simplifying studies on DNA replication (21). Fluorescence microscopy experiments showed that replisome components are diffuse in the cytoplasm of G1 swarmer cells and then assemble into a focus at the stalked pole of the cell at the onset of the replication process during the swarmer-to-stalked cell transition. As replication proceeds (S-phase), the replisome moves from the cell pole towards mid-cell, before it disassembles in late pre-divisional cells (22,23). *C. crescentus* finally divides into two different progenies: a swarmer G1-phase cell and a stalked S-phase cell. Thus, the sub-cellular localization of its moving replisome can be used as a *proxi* to visualize S-phase progression and analyze replication-associated processes. In this study, we characterized the MutH-independent MMR process of *C. crescentus* and its impact on genome maintenance, with a particular focus on its spatial coupling with DNA replication using fluorescently tagged MMR proteins and live cell fluorescence microscopy. It is the first detailed study on MMR in *Alphaproteobacteria*.

## MATERIALS AND METHODS

### Oligonucleotides, plasmids and strains

Oligonucleotides, plasmids and bacterial strains used in this study are listed in Tables S1, S2 and S3, respectively. Detailed methods used to construct plasmids and strains are also described in Supplementary Information.

### Growth conditions and synchronization procedure

*E. coli* strains were grown in Luria-Bertani (LB) broth or on LB + agar at 1.5% (LBA) at 37 °C. *C. crescentus* strains were cultivated at 28 °C in peptone yeast extract (PYE) medium or M2G minimal medium (24) with 180 rpm shaking or on PYE + agar 1.5% (PYEA). When needed, antibiotics were added to media (liquid/plates) at the following concentrations (μg/mL): gentamicin (15/20) or kanamycin (30/50) for *E. coli*; gentamicin (1/5), kanamycin (5/25), spectinomycin (25/100), rifampicin (−/5) for *C. crescentus*. When needed, xylose was used at a final concentration of 0.3% to induce the *Pxyl* promoter in *C. crescentus* (PYEX or M2GX). When needed, swarmer (G1-phase) cells were isolated from mixed populations of *C. crescentus* cells using a procedure adapted from (25): cells were first grown overnight in PYE medium and then diluted in M2G medium until cultures reached pre-exponential phase (OD_660nm_=0.1−0.2). Xylose 0.3% was added to induce the *xylX* promoter for 2.5 hours. The swarmer cells were then isolated by centrifugation in a Percoll density gradient and resuspended in M2G medium with 0.3% xylose.

### Spontaneous mutation frequency assays

The assay is based on spontaneous mutations that can occur in a specific region of the *rpoB* gene of *C. crescentus*, leading to rifampicin resistances; it was previously used as an efficient indicator of the spontaneous mutation rate to compare different strains (26). Here, we cultivated cells overnight in PYE medium and then diluted cultures into M2G medium to obtain a final 0.005<OD_660nm_<0.04. Growth was then continued overnight until cells reached exponential phase again (OD_660nm_=0.5). Serial dilutions of cultures were then prepared, and aliquots were plated onto PYEA with or without 5 μg/mL of rifampicin. To estimate the frequency at which spontaneous rifampicin resistant clones appeared in populations of cells, the number of rifampicin resistant colonies was divided by the total number of colonies that could grow without rifampicin. The assay was performed with minimum three independent cultures of each strain. Strains with genetic constructs expressed from the *xylX* promoter were cultivated in the presence of 0.3% xylose at all time during the procedure.

### Live cell microscopy

Two microscope systems were used to image cells during the course of this work: the first one is described in (27) and the second one in (28). Cells from strains compared to one another in a given figure were however always imaged using the same system. Cells were first cultivated overnight in PYE medium and then diluted in M2G medium to obtain a final 0.005<OD_660nm_<0.04. On the next day, once cultures reached early exponential phase (OD_660nm_~0.3), xylose 0.3% was added into the M2G medium when necessary to induce the *xylX* promoter. Once the culture reached an OD_660nm_ ~0.5, cells were immobilized onto a thin layer of M2 medium with 1% agarose on a slide prior to imaging.

For time lapse microscopy studies to follow swarmer (G1-phase) cells differentiating into stalked (early S-phase) and pre-divisional (late S-phase) cells, swarmer cells were isolated as described above (synchronization procedure) and immediately immobilized onto a thin layer of M2G medium complemented with 1% PYE, 0.3% xylose and 1% low melting temperature agarose (Promega) on slides. Slides were sealed (still with a significant air pocket) prior to imaging to prevent sample desiccation over time.

### Image analysis

Image analysis were performed using ImageJ and Photoshop to quantify the average fluorescent signal of the cytoplasm and the maximum fluorescent signal that could be detected in each cell. So-called “distinct foci” were arbitrarily defined if their maximum fluorescent signal was minimum two-fold higher than the average fluorescence intensity of the cytoplasm of each cell. Such experiments were performed minimum three times for each strain using independent cultures and images of >1500 cells were analysed. When needed, demographs were created with the Oufti software (29) using images of more than 700 cells for each culture using the parameters found in the file named “Caulobacter_crescentus_subpixel.set” (included in the software).

## RESULTS

### MutS, MutL and UvrD are critical for maintaining genome integrity in *C. crescentus*

A recent genetic screen looking for random *C. crescentus* mutator strains uncovered mutants with a transposon inserted into the *mutS* (*CCNA_00012*), *mutL* (*CCNA_00731*) or *uvrD* (*CCNA_01596*) genes (30). To confirm that they encode proteins involved in DNA repair, we constructed deletion mutants and compared their spontaneous mutation rate with that of an isogenic wild-type strain using classical rifampicin-resistance assays.

We found that Δ*mutS* and Δ*mutL* cultures formed spontaneous rifampicin-resistant colonies ~120-fold and ~110-fold more frequently than wild-type cultures (Fig.1 and Table S4). Importantly, a double mutant carrying both mutations displayed a mutation rate only slightly elevated (~139-fold more than the wild-type strain) compared to single mutants (Fig.1 and Table S4), providing a strong indication that MutS and MutL mostly function in the same MMR pathway, as expected. Notably, amino-acid residues required for mismatch recognition and nucleotide binding/hydrolysis by the *E. coli* and *B. subtilis* MutS proteins (Fig.S1), as well as amino-acid residues required for the endonuclease activity of the *B. subtilis* MutL (Fig.S2), were found to be conserved on *C. crescentus* MutS and MutL. Thus, it is very likely that MutS detects DNA mismatches, while MutL cleaves the DNA strand that needs to be repaired during the *C. crescentus* MMR pathway, as it is the case in *B. subtilis*.

**Figure 1:**
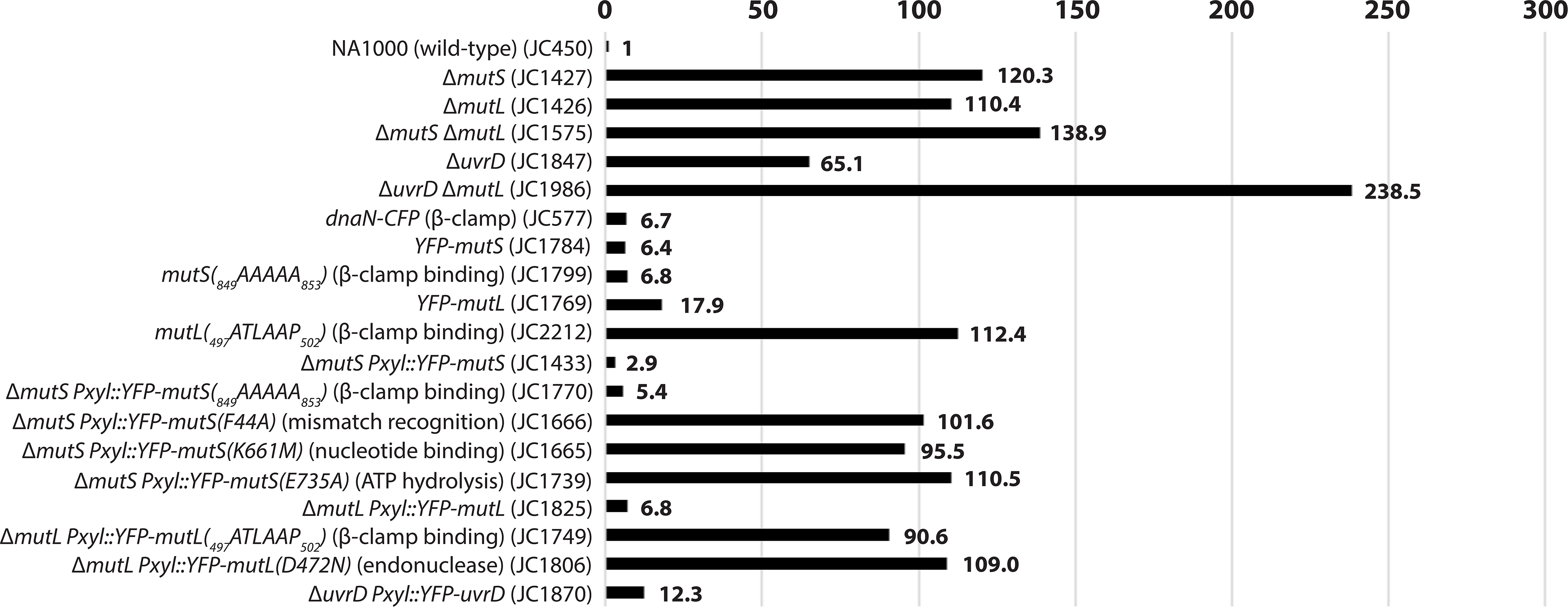
Comparison of the spontaneous mutation rates of different *C. crescentus* strains. This figure is based on values described in Table S4. Relevant genotypes (and strain numbers) are indicated on the left side of the figure. To facilitate comparisons, values were normalized so that the value for a wild-type NA1000 strain equals 1. The spontaneous mutation rate of each strain was estimated by measuring the spontaneous appearance of rifampicin-resistant clones. Each value was estimated from minimum three independent cultures (standard deviations are described in Table S4).

Δ*uvrD* cultures formed ~65-fold more rifampicin-resistant colonies than wild-type cultures (Fig.1 and Table S4), also consistent with a role of the *C. crescentus* UvrD in DNA repair. However, a double mutant lacking *mutL* and *uvrD* displayed a higher mutation rate than the corresponding single mutants (~238-fold more rifampicin-resistant colonies than wild-type for the double mutant, compared to ~110- or ~65-fold more for single mutants) (Fig.1 and Table S4). This observation provided a first indication that UvrD is involved in minimum one other DNA repair pathway(s) beyond MMR in *C. crescentus*: it is most likely the nucleotide excision repair (NER) pathway mediated by UvrABC in bacteria (30).

### MutS co-localizes with the replisome throughout the S-phase of the *C. crescentus* cell cycle

Considering that DNA mismatches are mostly generated by the mis-incorporation of nucleotides by the replicative DNA polymerase and that previous studies on other MutS homologs had shown that they are sometimes associated with the replisome in bacterial cells (8,15), we looked at the sub-cellular localization of MutS in *C. crescentus*. To get started, we constructed a strain expressing a fluorescently tagged YFP-MutS protein from the native *mutS* promoter and replacing MutS. Importantly, we found that the spontaneous mutation rate of this strain was very close to that of the wild-type strain (Fig.1 and Table S4), demonstrating that YFP-MutS is almost fully functional. Unfortunately, the fluorescence signal displayed by these cells was too low to be detected using our microscopy setups (data not shown). Then, we constructed another strain expressing YFP-MutS from the chromosomal xylose-inducible *xylX* promoter (*Pxyl*) in an otherwise Δ*mutS* background. Similar to the *YFP-mutS* strain, the MMR process appeared as almost fully functional (~98% of activity) in this Δ*mutS Pxyl::YFP-mutS* strain (Fig.1 and Table S4). Once we had checked by immunoblotting that the YFP moiety of YFP-MutS remained bound to MutS *in vivo* (Fig. S3), we proceeded with live cells fluorescence microscopy experiments. We observed that the fluorescent signal was essentially spread throughout the cytoplasm of ~30% of the cells, while it formed distinct foci (signal >2-fold above the cytoplasmic signal) in ~70% of the cells (Fig. 2A). When detectable, foci localized at one cell pole or at a position between the cell pole and mid-cell. Remarkably, a classification of cells according to their size (Fig.3A) showed that the shortest (swarmer/G1) cells usually displayed no focus, that longer (stalked/early S-phase) cells displayed foci close to the cell pole, while even longer (early pre-divisional/late S-phase) cells displayed foci near mid-cell. A time-lapse microscopy experiment following the cell cycle of newly born swarmer/G1 cells (Fig.3B) confirmed that the sub-cellular localization of MutS was very similar to that of the replisome (22,23,31) (Fig.3C).

**Figure 2:**
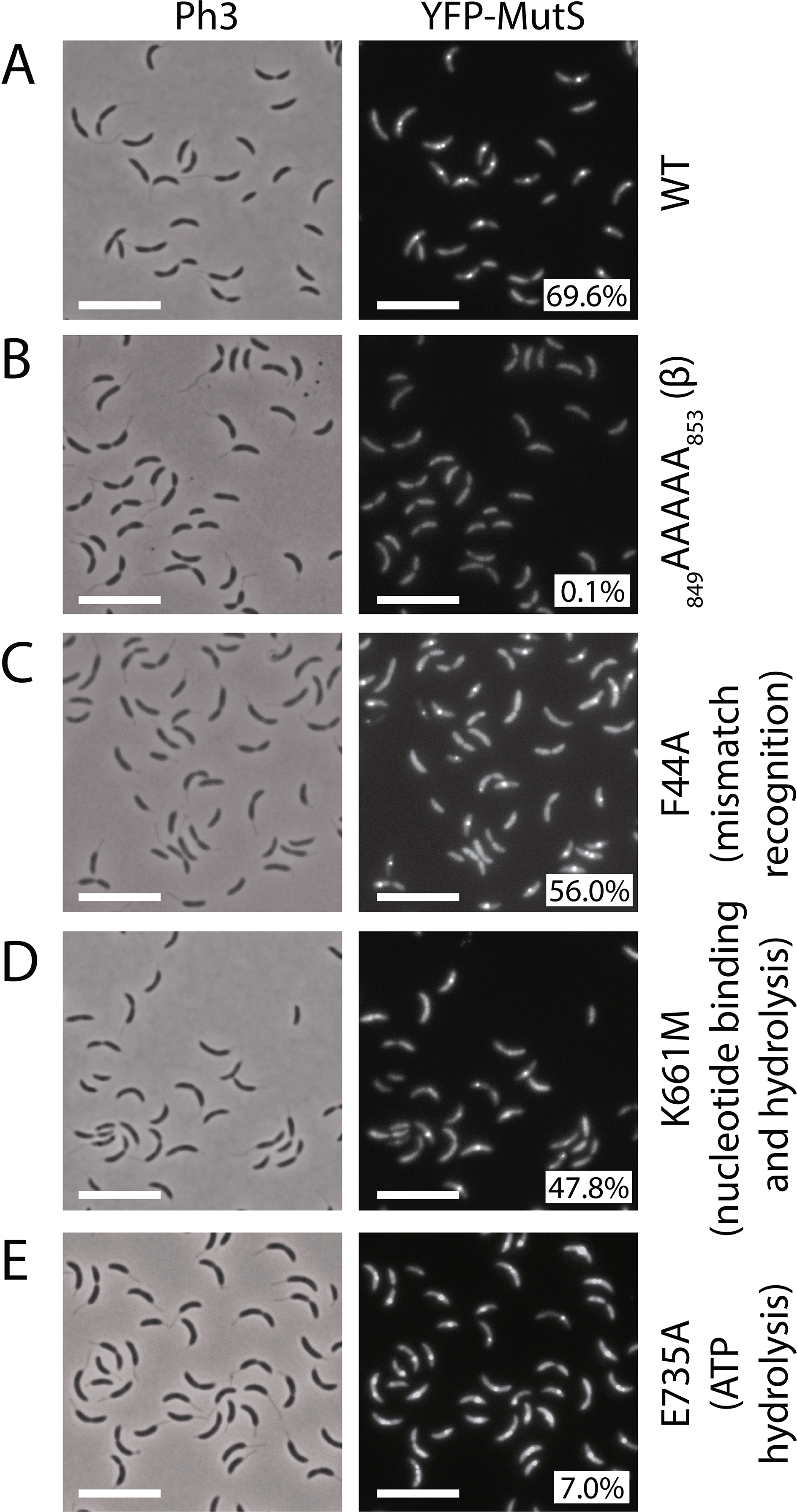
YFP-MutS forms discrete fluorescent foci in a majority of *C. crescentus* cells. The subcellular localization of several derivatives of YFP-MutS was analyzed in Δ*mutS* cells. Strains JC1433 (Δ*mutS Pxyl::YFP-mutS*) **(A)**, JC1770 (Δ*mutS Pxyl::YFP-mutS(849AAAAA853)*) **(B)**, JC1666 (Δ*mutS Pxyl::YFP-mutS(F44A)*) **(C)**, JC1665 (Δ*mutS Pxyl::YFP-mutS(K661M)*) **(D)** and JC1739 (Δ*mutS Pxyl::YFP-mutS(E735A)*) **(E)** were cultivated into PYE medium and then transferred into M2G medium. 0.3% xylose was added to cultures when they reached an OD_660nm_~0.3. Cells were then imaged by fluorescence microscopy when the OD_660nm_ reached ~0.5. Representative images are shown here. Ph3 indicates phase-contrast images. The % indicated onto images corresponds to the average proportion of cells (using values obtained from three independent experiments) displaying a distinct fluorescent focus (intensity >2-fold above background). The white scale bar corresponds to 8μm.

**Figure 3:**
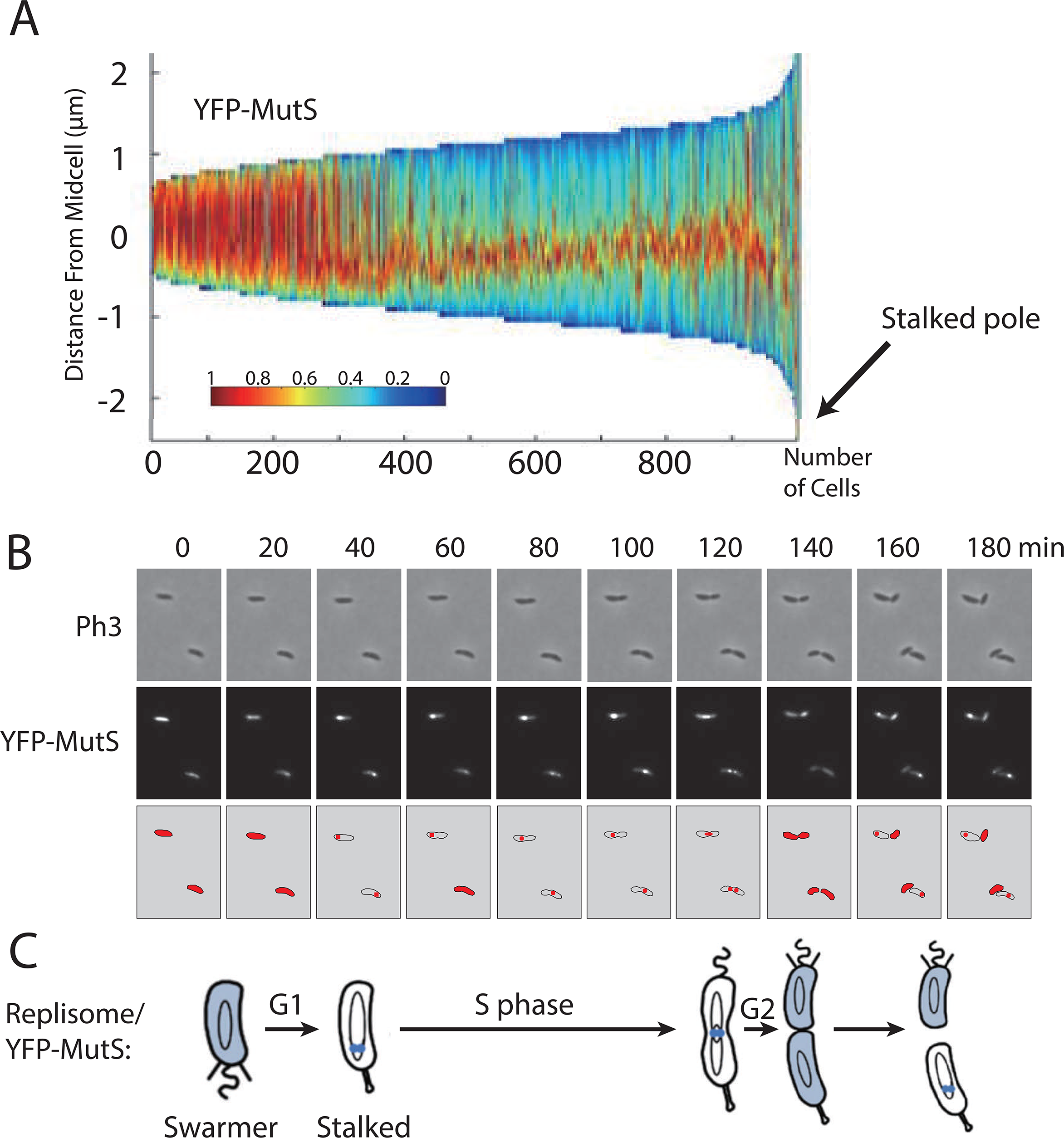
YFP-MutS forms discrete fluorescent foci throughout the S-phase of the *C. crescentus* cell cycle. **(A)** Demograph showing the subcellular localization of YFP-MutS in Δ*mutS* cells sorted as a function of their size. JC1433 (Δ*mutS Pxyl::YFP-mutS*) cells were cultivated and imaged as described for Fig.2. Short cells correspond to G1/swarmer cells, while intermediate and longer cells correspond to stalked and pre-divisional S-phase cells, respectively. **(B)** Time-lapse fluorescence microscopy experiment showing the cell cycle localization of YFP-MutS as a function of the cell cycle of Δ*mutS* cells. JC1433 cells were first cultivated in PYE medium overnight and then diluted in M2G medium until the cells reached pre-exponential phase (OD_660nm_=0.1-0.2). Xylose at 0.3% was added into the cultures to induce the *Pxyl* promoter for 2.5 hours. Swarmer cells were then isolated (synchronization protocol) from the cell culture, immobilized onto an agarose pad and imaged by fluorescence microscopy every 20 minutes. Representative images are shown here. The schematics drawn under the microscopy images highlight in red the subcellular localization of YFP-MutS in cells imaged above. **(C)** This schematic shows the *C. crescentus* cell cycle and the blue color highlights where MutS appears to be localized as a function of the cell cycle using results from panels A and B. This localization pattern is reminiscent of the known localization pattern of replisome components in *C. crescentus* (22).

To show more directly that YFP-MutS foci are co-localized with the replisome, we introduced a *dnaN-CFP* construct replacing the native *dnaN* gene (27) into the Δ*mutS Pxyl::YFP-mutS* strain. The *dnaN-CFP* allele had an only very minor impact on the mutation rate of strains carrying it (Fig.1 and Table S4), but we found unexpectedly that the CFP moiety added to DnaN disturbed the proportion of cells displaying YFP-MutS foci (~22% instead of ~70% of cells with distinct YFP-MutS foci) (Fig. S4). It still proved somewhat informative to show that the vast majority (~96%) of these distinct YFP-MutS foci were co-localized with the DnaN β-clamp of the DNA polymerase.

If an association between MutS and the replisome is responsible for the particular localization pattern of MutS, one would also expect that focus formation would be disturbed or inhibited in non-replicating cells. To test this more directly, we treated cells expressing YFP-MutS with novobiocin, a drug that inhibits the DNA gyrase and leads to replisome disassembly in *C. crescentus* (22,23): only very few cells (1%) still exhibited distinct YFP-MutS foci by fluorescence microscopy (Fig. S5), indicating that ongoing replication is required for YFP-MutS foci formation/maintenance.

Altogether, these results revealed that MutS associates with the replisome in a rather stable manner throughout the whole S-phase of the *C. crescentus* cell cycle.

### The putative β-clamp binding motif of MutS is critical for its recruitment at the replisome, but not for its activity in *C. crescentus*

The *C. crescentus* MutS protein carries a motif (_849_DLPLF_853_) close to its C-terminus, which shows some similarities with the β-clamp binding motifs of the *E. coli* (_812_QMSLL_816_) and *B. subtilis* (_806_QLSFF_810_) MutS proteins (Fig.S1). To test if this motif is involved in the recruitment of MutS to the replisome in *C. crescentus*, we constructed a Δ*mutS* strain expressing a mutant YFP-MutS(_849_AAAAA_853_) protein from the *Pxyl* promoter. As predicted, fewer than 0.2% of the cells expressing this mutant protein displayed distinct YFP foci (Fig.2B), even when expressed in *dnaN-CFP* cells that displayed frequent CFP foci (distinct in ~54% of cells) (Fig.S4). These observations show that the _849_DLPLF_853_ β-clamp binding motif of MutS is necessary for the spatial association between MutS and the replisome in *C. crescentus* cells, strongly suggesting that the β-clamp recruits MutS to the replisome during the S-phase of the cell cycle.

We next wished to use this mutation to test if the spatial association between MutS and the replisome is necessary or useful during the *C. crescentus* MMR process. We compared the spontaneous mutation rates of Δ*mutS* strains expressing either YFP-MutS or YFP-MutS(_849_AAAAA_853_) from the *xylX* promoter and found similar rates (Fig.1 and Table S4). As a second check, we also replaced the native *mutS* allele of a wild-type strain with the mutant *mutS(_849_AAAAA_853_)* allele for expression at native levels and still did not observe obvious differences in mutation rates (Fig.1 and Table S4). Thus, we conclude that the spatial association of MutS with the replisome is strong during the S-phase of the cell cycle, but not necessary for the MMR process, at least under classical laboratory growth conditions that do not promote mismatch occurrence.

### Mismatch frequency, or the capacity of MutS to detect mismatches, do not affect MutS localization in *C. crescentus*

To test if the localization of MutS was influenced by the frequency at which mismatches occur in *C. crescentus*, we constructed a novel mutator strain with a *dnaQ(G13E)* allele replacing the native *dnaQ* (*CCNA_00005*) gene. Since DnaQ epsilon sub-units of bacterial DNA polymerases III carry their proofreading activity (32), the spontaneous mutation rate of a *C. crescentus dnaQ(G13E)* strain was largely increased (~479-fold) compared to the wild-type strain (Table S4). This mutation was then introduced into the Δ*mutS Pxyl::yfp-mutS* strain for fluorescence microscopy experiments. Interestingly, the proportion of cells displaying YFP-MutS foci was essentially identical in wild-type and *dnaQ(G13E)* cells (Fig.4). Thus, the subcellular localization of MutS does not appear to be influenced by the frequency of mismatches in *C. crescentus* cells.

**Figure 4:**
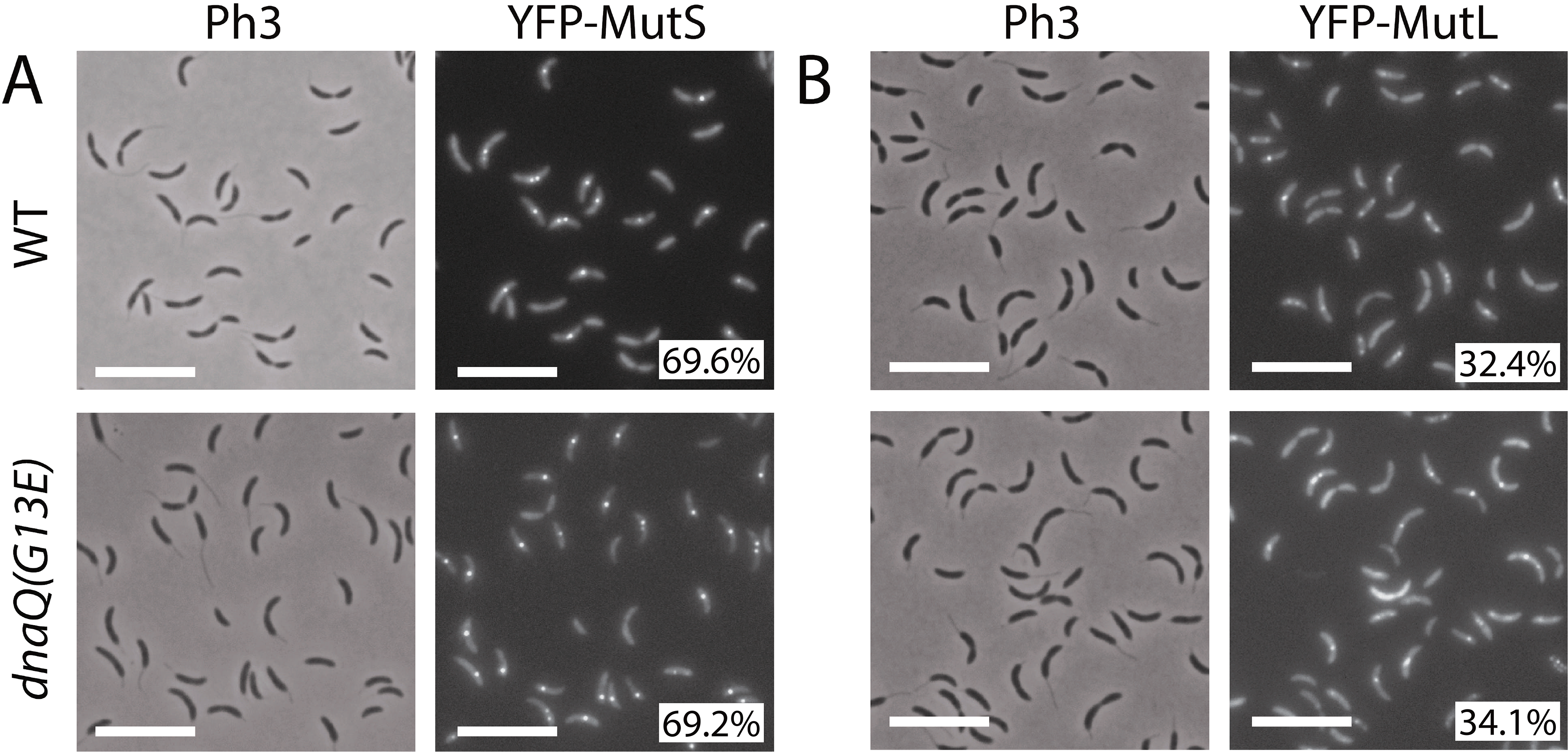
YFP-MutS and YFP-MutL form frequent foci in *C. crescentus* cells, independently of mismatch frequency. **(A)** YFP-MutS localization in cells with a wild-type (WT) or a proofreading-deficient (*dnaQ(G13E)*) replicative DNA polymerase. Strains JC1433 (Δ*mutS Pxyl::YFP-mutS*) and JC1724 (Δ*mutS Pxyl::YFP-mutS dnaQ(G13E)*) were cultivated into PYE medium and then transferred into M2G medium. 0.3% xylose was added to cultures when they reached an OD_660nm_~0.3. Cells were then imaged by fluorescence microscopy when the OD_660nm_ reached ~0.5. **(B)** YFP-MutL localization in cells with a wild-type or a proofreading-deficient replicative DNA polymerase. Strains JC1825 (Δ*mutL Pxyl::YFP-mutL*), and JC1845 (Δ*mutL Pxyl::YFP-mutL dnaQ(G13E)*) were cultivated and imaged as described for panel A. Representative images are shown in panels A and B. Ph3 indicates phase-contrast images. The % indicated onto images corresponds to the average proportion of cells (using values obtained from three independent experiments) displaying a distinct fluorescent focus (intensity >2-fold above background). The white scale bar corresponds to 8μm.

To confirm that mismatch detection by MutS is not a pre-requisite for the recruitment of MutS to the replisome, we also characterized the localization of a mutant YFP-MutS(F44A) protein that carries a point mutation in its predicted mismatch detection motif (_42_<u>G</u>D<u>FY</u>ELFFD<u>DA</u>_52_ in Fig.S1). As expected, *ΔmutS* cultures expressing *YFP-mutS(F44A)* generated nearly as many spontaneous mutations as Δ*mutS* cultures (Fig.1 and Table S4), showing that MutS(F44A) is mostly non-functional. Still, YFP-MutS(F44A) formed distinct fluorescent foci in ~56% of *ΔmutS* cells (Fig.2C), showing that efficient mismatch detection by MutS is not a pre-requisite for focus formation. Moreover, Δ*mutS dnaN-CFP* cells expressing *YFP-mutS(F44A)* displayed YFP-MutS foci as frequently as isogenic cells expressing *YFP-mutS*, and these foci were still co-localized with DnaN-CFP foci (Fig. S4).

Altogether, these results indicate that the spatial coupling between MutS and the replisome is essentially independent of mismatch recognition by MutS in *C. crescentus*.

### The nucleotide binding to MutS contributes to its activity and affects its localization at the replisome in *C. crescentus*

To gain insight into the impact of nucleotide binding/hydrolysis on MutS activity and localization, the two predicted Walker motifs of the *C. crescentus* MutS protein (Fig.S1) were mutagenized. The Walker A motif was disrupted in the YFP-MutS(K661M) protein and the Walker B motif was disrupted in the YFP-MutS(E735A) protein. Cultures of cells expressing these mutant *mutS* alleles as the sole copy of *mutS* displayed a much higher mutation rate (33-fold and 38-fold, respectively) than cells expressing the wild-type *mutS* allele at similar levels (Fig.1 and Table S4), indicating that ATP binding and/or hydrolysis on MutS is important during the *C. crescentus* MMR process. Moreover, comparison of the two mutants suggests that MutS bound to ATP may be severely impaired in its capacity to detect mismatches (92% loss of activity for YFP-MutS(E735A) that can supposedly not hydrolyse ATP), while unbound MutS may still keep some activity (79% loss of activity for YFP-MutS(K661M) that can supposedly not bind to ATP/ADP).

We then characterized the sub-cellular localization of these mutant proteins by fluorescence microscopy. Using the Δ*mutS Pxyl::YFP-mutS(K661M)* strain, we found that YFP-MutS(K661M) formed frequent foci, but less frequently than YFP-MutS (~48%, instead of ~70% of cells with a distinct focus) (Fig.2D). Furthermore, observation of Δ*mutS dnaN-CFP Pxyl::YFP-mutS(K661M)* cells showed that YFP-MutS(K661M) formed only very rare replisome-associated foci in replicating cells (~3%) (Fig.S4). Microscopy using Δ*mutS Pxyl::YFP-mutS(E735A)* cells showed that YFP-MutS(E735A) formed rare distinct foci (~7%) (Fig.2E), while microscopy using Δ*mutS dnaN-CFP Pxyl::YFP-mutS(E735A)* cells showed that YFP-MutS(E735A) formed rare replisome-associated foci in replicating cells (~8%) (Fig.S4). Overall, these observations suggest that unbound MutS may have less affinity for the replisome than ADP-bound MutS, while ATP-bound MutS may display the lowest affinity, which is reminiscent of the so-called “sliding clamp” model following mismatch detection by MutS in *C. crescentus.*

### YFP-MutL forms replisome-associated foci in a subset of S-phase *C. crescentus* cells and independently of mismatch formation

Knowing that MutS is found associated with the replisome during the S-phase of the cell cycle, the sub-cellular localization of MutL was also analyzed. As we did for *mutS*, we first replaced the native wild-type allele of *mutL* with a *yfp-mutL* allele expressed from the native *mutL* promoter on the *C. crescentus* chromosome. This strain displayed a spontaneous mutation rate quite similar to the wild-type strain (Fig.1 and Table S4), indicating that YFP-MutL can still repair ~85% of the mismatches that normally get repaired by MutL. The fluorescence signal displayed by these cells was unfortunately too low to be detected using our microscopy setups (data not shown). Then, we switched to a Δ*mutL* strain expressing *YFP-mutL* from the chromosomal *Pxyl* promoter for subsequent fluorescence microscopy analysis. This strain displayed a spontaneous mutation rate even closer to the wild-type strain than the *YFP-mutL* strain (Fig.1 and Table S4): we estimated that YFP-MutL expressed from the *Pxyl* promoter corrects ~95% of the mismatches corrected by MutL expressed from the native *mutL* promoter, confirming that YFP-MutL is almost fully functional. Immunoblotting experiments also showed that the YFP moiety remained bound to YFP-MutL *in vivo* (Fig.S3). Analysis of these cells by fluorescence microscopy revealed that ~32% of the cells displayed distinct fluorescent foci (with a signal >2-fold above the cytoplasmic signal) (Fig.5A), a number significantly lower than previously found with Δ*mutS Pxyl::yfp-mutS* cells (~70%) (Fig.2A). Still, these YFP-MutL foci appeared as dependent on ongoing replication like YFP-MutS foci, since they largely disassembled following a novobiocin treatment (Fig.S5), while at the same time being more transient than YFP-MutS foci, since we observed by time-lapse microscopy (Fig.5B) that single YFP-MutL foci could assemble (example at time point 80’ in Fig.5B) and disassemble (example at time point 100’ in Fig.5B) within the same cell cycle. Interestingly, we also observed that YFP-MutL foci were nearly never detected in short swarmer cells, only rarely detected in stalked cells and more often detected in pre-divisional cells (Fig.5B and Fig.S6). Altogether, these first observations suggested that YFP-MutL may not associate with the replisome as frequently or with as much affinity as YFP-MutS, especially at the beginning of the S-phase.

**Figure 5:**
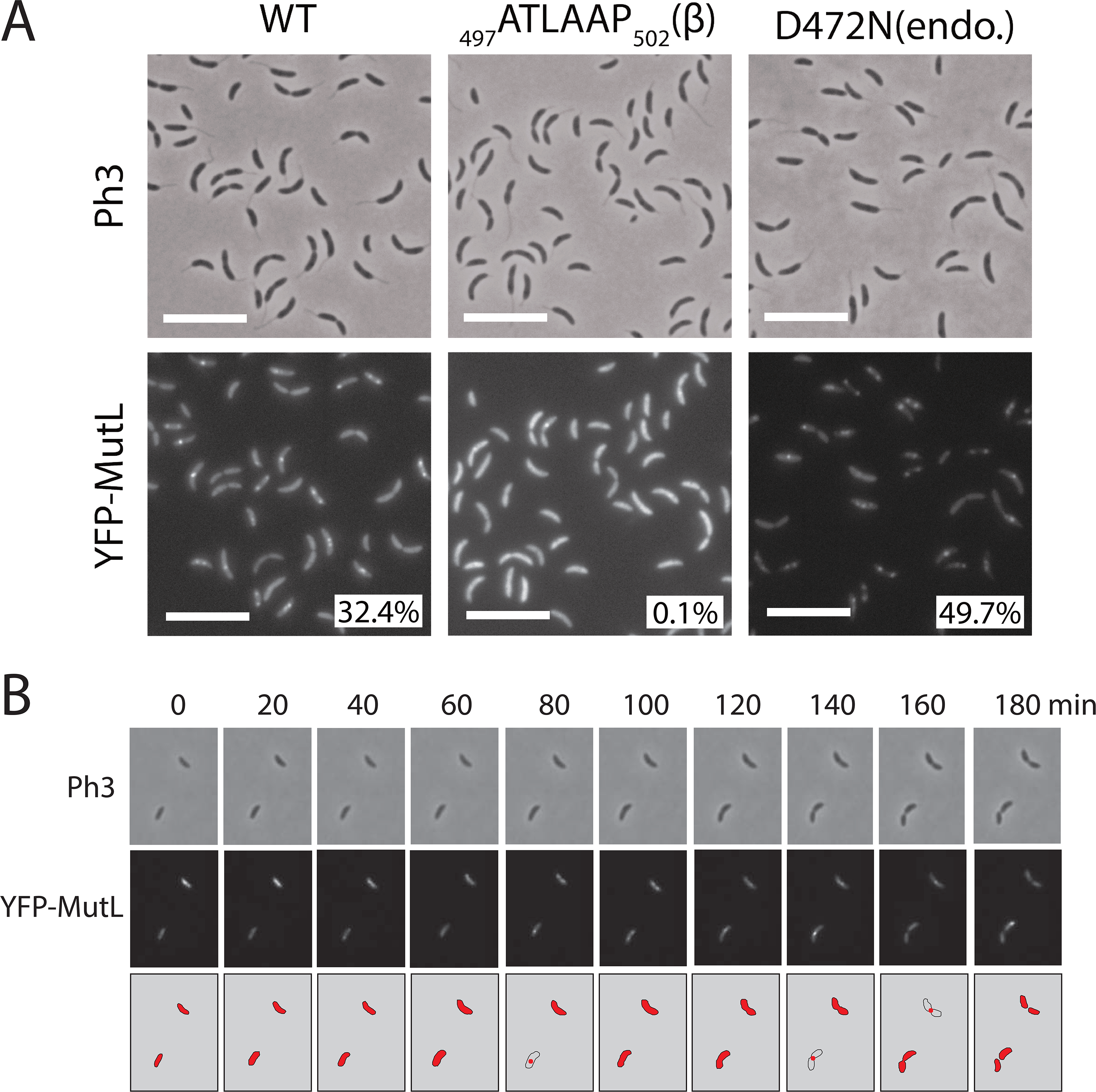
YFP-MutL forms discrete fluorescent foci in a subset of S-phase *C. crescentus* cells. **(A)** Subcellular localization of several derivatives of YFP-MutL in Δ*mutL* cells. Strains JC1825 (Δ*mutL Pxyl::YFP-mutL*) labelled “WT”, JC1749 (Δ*mutL Pxyl::YFP-mutL(_497_ATLAAP_502_)*) labelled “β" and JC1667 (Δ*mutL Pxyl::YFP-mutL(D472N))* labelled “endo” were cultivated into PYE medium and then transferred into M2G medium. 0.3% xylose was added to cultures when they reached an OD_660nm_~0.3. Cells were then imaged by fluorescence microscopy when the OD_660nm_ reached ~0.5. The % indicated onto images corresponds to the average proportion of cells (using values obtained from three independent experiments) displaying a distinct fluorescent focus (intensity >2-fold above background). The white scale bar corresponds to 8μm. **(B)** Time-lapse fluorescence microscopy experiment showing the localization of YFP-MutL as a function of the cell cycle of Δ*mutL* cells. JC1825 cells were first cultivated in PYE medium overnight and then diluted in M2G medium until the cells reached pre-exponential phase (OD_660nm_=0.1-0.2). Xylose at 0.3% was added into the cultures to induce the *Pxyl* promoter for 2.5 hours. Swarmer cells were then isolated (synchronization protocol) from the cell culture, immobilized onto an agarose pad and imaged by fluorescence microscopy every 20 minutes. The schematics drawn under the microscopy images highlight the localization of YFP-MutL in cells imaged above. Representative images of cells are shown and Ph3 indicates phase-contrast images in panels A and B.

To shed light on this rather dynamic localization pattern for MutL, we carefully analyzed the localization of DnaN-CFP and YFP-MutL in cells expressing both proteins simultaneously. We found that ~97% of the YFP-MutL foci that can be detected are co-localized with DnaN-CFP foci (Fig.6A), showing that YFP-MutL is nearly always associated with the replisome when it forms foci. Sorting cells as a function of their size also confirmed that YFP-MutL foci are mainly detected during the S-phase of the cell cycle, being apparently more often associated with the replisome towards the end of the S-phase (Fig.6B&C).

**Figure 6:**
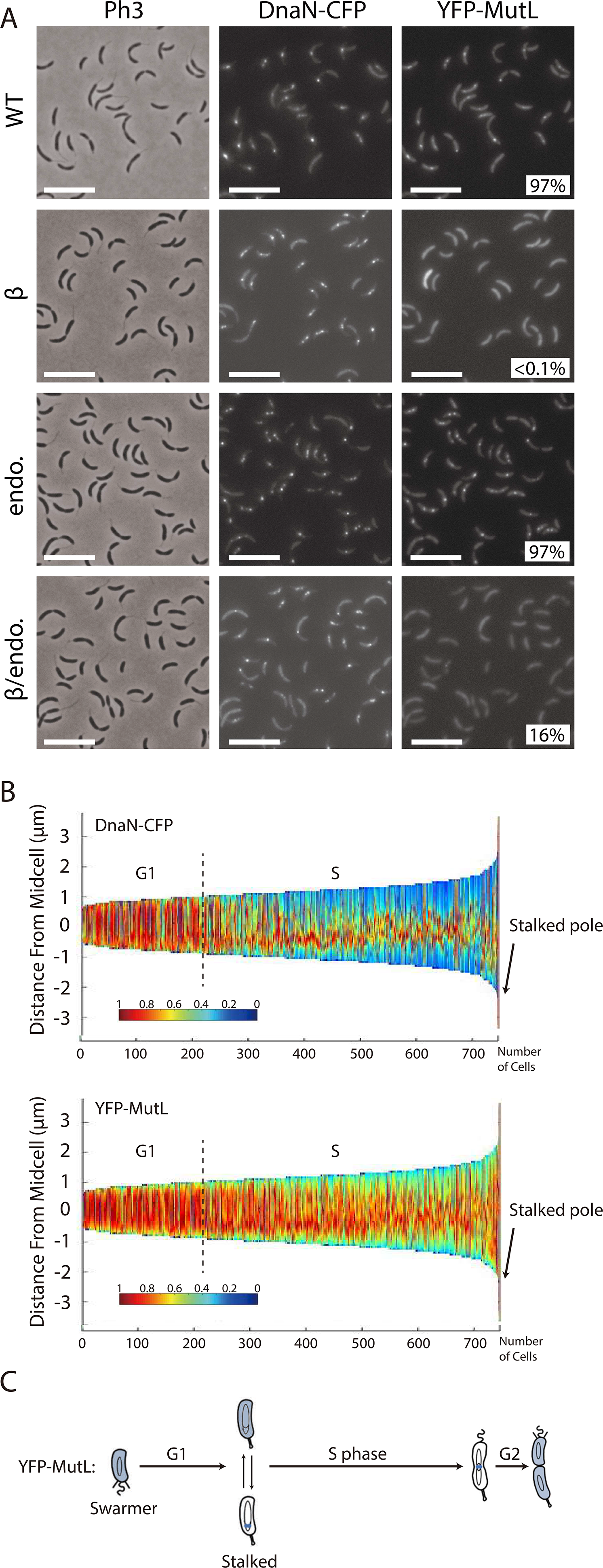
YFP-MutL foci co-localize with the replisome. **(A)** Subcellular localization of DnaN-CFP and of several derivatives of YFP-MutL in Δ*mutL* cells. Strains JC1812 (*dnaN-CFP* Δ*mutL Pxyl::YFP-mutL*) labelled “WT”, JC1750 (*dnaN-CFP* Δ*mutL Pxyl::YFP-mutL(_497_ATLAAP_502_)*) labelled “β", JC1670 (*dnaN-CFP* Δ*mutL Pxyl::YFP-mutL(D472N)*) labelled “endo” and JC1753 (*dnaN-CFP* Δ*mutL Pxyl::YFP-mutL(D472N-497ATLAAP502)*) labelled “β/endo”, were cultivated into PYE medium and then transferred into M2G medium. 0.3% xylose was added to cultures when they reached an OD_660nm_~0.3. Cells were then imaged by fluorescence microscopy when the OD_660nm_ reached ~0.5. The % indicated onto images corresponds to the average proportion of distinct MutL-YFP foci (intensity >2-fold above average background) that are co-localized with DnaN-CFP foci (using values obtained from three independent experiments). The white scale bar corresponds to 8μm. **(B)** Demographs showing the subcellular localization of DnaN-CFP and YFP-MutL in Δ*mutL* cells sorted as a function of their size. Strain JC1812 was cultivated and imaged as described for panel A. Short cells correspond to G1/swarmer cells, while intermediate and longer cells correspond to stalked and pre-divisional S-phase cells, respectively. **(C)** This schematic shows the *C. crescentus* cell cycle and the blue color highlights where YFP-MutL is localized as a function of the cell cycle based on images shown in panel B and in Fig.5B.

As observed for YFP-MutS foci, these replisome-associated YFP-MutL foci appeared as independent of the frequency of mismatch occurrence, since the introduction of the *dnaQ(G31E)* allele in this strain did not affect the localization pattern or the proportion of cells displaying YFP-MutL foci (Fig.4B). Consistently, the association of MutL with the replisome did not require mismatch detection by MutS, since YFP-MutL foci were still co-localized with DnaN-CFP in *ΔmutS* cells (Fig.S7).

### MutL is recruited to the replisome through a putative β-clamp binding motif that is critical for MMR in *C. crescentus*

Considering that MutL can associate with the replisome in the absence of MutS in *C. crescentus* (Fig.S7), and that MutL binds directly to the β-clamp of the DNA polymerase in other bacterial species (11,15), we searched for a putative β-clamp binding motif on the *C. crescentus* MutL protein. We found a _497_QTLLLP_502_ motif (Fig. S2) closely related with the previously proposed Qxh(L/I)xP consensus β-clamp binding motif of MutL proteins (33). We therefore engineered a *Pxyl::YFP-mutL(_497_ATLAAP_502_)* construct and introduced it into *C. crescentus* Δ*mutL* and Δ*mutL dnaN-CFP* strains. We imaged cells from both strains by fluorescence microscopy and found that fewer than 0.1% of the cells displayed a distinct YFP-MutL(_497_ATLAAP_502_) focus (Fig.5A and Fig.6A). Moreover, none of the replicating cells from the second strain (~60% of the cells that displayed DnaN-CFP foci) displayed a YFP-MutL(_497_ATLAAP_502_) focus that co-localized with a DnaN-CFP focus (Fig. 6A). This finding shows that the _497_QTLLLP_502_ motif of MutL is critical for focus formation and it is a strong indication that MutL is recruited to the replisome through a direct interaction with the β-clamp.

We then estimated the spontaneous mutation rate of the *ΔmutL Pxyl::YFP-mutL(_497_ATLAAP_502_)* strain to test if the recruitment of MutL to the replisome contributes to MutL activity (Fig.1 and Table S4). We found that this strain made ~13-fold more mutations than the isogenic strain expressing the YFP-MutL protein at similar levels (Fig.1 and Table S4), indicating that YFP-MutL(_497_ATLAAP_502_) (expressed form the *Pxyl* promoter) is mostly inactive. To verify this result when *mutL* is expressed from its native promoter and in the absence of the *yfp* moiety, we also replaced the native *mutL* gene by the mutant *mutL(_497_ATLAAP_502_)* allele on the *C. crescentus* chromosome. Strikingly, the spontaneous mutation rate of this *mutL(_497_ATLAAP_502_)* strain was essentially identical to that of a Δ*mutL* strain (~112-fold higher than the wild-type strain) (Fig.1 and Table S4), showing that MutL(_497_ATLAAP_502_) is totally inactive. All together, these results suggest that the MutS- and mismatch-independent recruitment of MutL to the replisome may licence the endonuclease activity of MutL, which is predicted to be the essential activity of MutL during the MMR process in *C. crescentus*.

### An inactive MutL(D472N) protein is stabilized at the replisome in *C. crescentus*

To gain insight into the connection between MutL recruitment to the replisome and its activity as an endonuclease during the MMR process, we engineered a mutant YFP-MutL(D472N) protein that lacks the conserved aspartate residue in its predicted endonuclease domain (34) (Fig.S2). As expected, a strain expressing *YFP-MutL(D472N)* as the only copy of *mutL* on the chromosome has a mutation rate nearly identical to that of a Δ*mutL* strain (Fig.1 and Table S4), demonstrating that MutL(D472N) is completely inactive for MMR. Interestingly, we observed by fluorescence microscopy that YFP-MutL(D472N) formed foci significantly more frequently than YFP-MutL: ~50% instead of ~32% of the cells displayed distinct foci (>2-fold above cytoplasmic signal) (Fig.5A). Using a *dnaN-CFP* derivative of that strain, we found that ~83% of the S-phase cells displayed YFP-MutL(D472N) foci that co-localized with DnaN-CFP foci, which was significantly higher than what was observed for YFP-MutL (~58%) (Fig.6A). Thus, MutL appears to be stabilized at the replisome when it loses its endonuclease activity. Importantly, YFP-MutL(D472N) was still frequently associated with the replisome in Δ*mutS* cells (Fig. S8), confirming that MutL recruitment to the replisome is independent of mismatch detection by MutS.

We also tested whether the stabilization of MutL(D472N) at the replisome was dependent on its β-clamp binding motif. Microscopy analysis of Δ*mutL dnaN-CFP Pxyl::YFP-mutL(D472N, _497_ATLAAP_502_)* cells showed that only ~1% of the replicating cells (with a DnaN-CFP focus) displayed a YFP-MutL(D472N, _497_ATLAAP_502_) focus that co-localized with the DnaN-CFP focus, which was dramatically lower than what was observed using isogenic cells expressing YFP-MutL(D472N) instead (~83%) (Fig.6A). Then, YFP-MutL(D472N) is stabilized at the replisome in a manner that is dependent on its interaction with the β-clamp.

Altogether, our results on the *C. crescentus* MutL protein suggest that it is active as an endonuclease when it is at the replisome and that this activity also influences its dynamic association with the replisome.

### YFP-UvrD forms rare and mostly MutS- and mismatch-independent foci in *C. crescentus*

To gain insight on whether the UvrD helicase may play a role during the MMR process in *C. crescentus*, as it is the case in *E. coli* (35), we also characterized its sub-cellular localization in *C. crescentus* cells. We constructed a Δ*uvrD* strain expressing a fluorescently tagged YFP-UvrD protein from the chromosomal *Pxyl* promoter. This strain displayed a spontaneous mutation rate slightly but significantly higher than that of a wild-type strain, suggesting that YFP-UvrD retains ~82% of its activity (Fig.1 and Table S4). We also checked by immunoblotting that its YFP moiety remained bound to UvrD *in vivo* (Fig.S3) prior to imaging cells by fluorescence microscopy. We found that only ~3% of *Pxyl::YFP-uvrD* cells displayed YFP-UvrD foci (with a signal >2-fold above the cytoplasmic signal) (Fig.7). These rare foci were found at any position in the cytoplasm of cells. In order to test if these foci may be connected with the repair of mismatches generated by the replicative DNA polymerase, we looked at the influence of mismatch occurrence on the assembly of YFP-UvrD foci. To do so, we introduced the *dnaQ(G13E)* mutation into these cells. Fluorescence microscopy analysis using these cells showed the *dnaQ(G13E)* mutation does not influence the proportion of cells displaying YFP-UvrD foci (Fig.7), suggesting that they may not correspond to active MMR sites. Consistent with this proposal, we also found that YFP-UvrD foci assembled nearly as frequently in Δ*mutS* than in wild-type cells (~2% versus ~3%, respectively) (Fig.7). These microscopy observations, together with the comparison of the spontaneous mutation rates of single and double mutants of *uvrD* and/or *mutL* (Fig.1 and Table S4), suggest that the main function of UvrD in maintaining genome integrity is not solely (or not at all) through a contribution to the MMR process. Instead, most of the YFP-UvrD foci may represent active NER sites.

**Figure 7:**
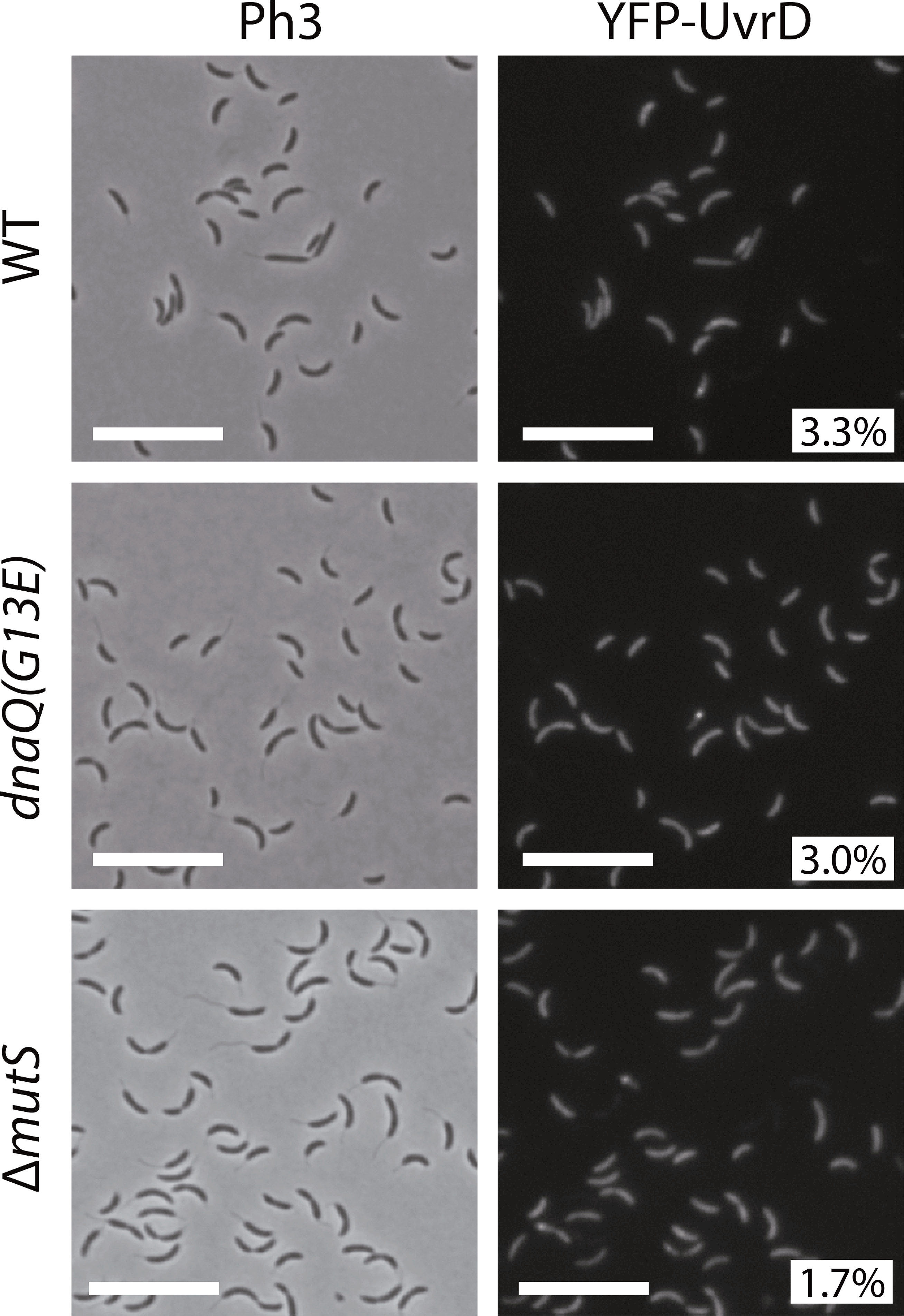
YFP-UvrD forms rare fluorescent foci in *C. crescentus* cells. Subcellular localization of YFP-UvrD in wild-type*, dnaQ(G13E)* or Δ*mutS* cells. Strains JC1946 (*Pxyl::YFP-uvrD*), JC2211 (*dnaQ(G13E) Pxyl::YFP-uvrD*) and JC1977 (Δ*mutS Pxyl::YFP-uvrD*) were cultivated into PYE medium and then transferred into M2G medium. 0.3% xylose was added to cultures when they reached an OD_660nm_~0.3. Cells were then imaged by fluorescence microscopy when the OD_660nm_ reached ~0.5. Representative images are shown here. Ph3 indicates phase-contrast images. The % indicated onto images corresponds to the average proportion of cells (using values obtained from three independent experiments) displaying a fluorescent focus (intensity >2-fold above background). The white scale bar corresponds to 11μm.

## DISCUSSION

In this study, we estimated that ~99.8% of the bases that are incorrectly incorporated by the DNA polymerase III of *C. crescentus* are detected and removed by its DnaQ-dependent proofreading activity (Table S4). Still, a significant number of mismatches escape this control system and must be removed before they turn into permanent mutations to ensure genome stability over generations. Here, we found that the *C. crescentus* MMR system is spatially associated with the replisome to detect and then correct ~99% of these left-over mismatches (Fig.1 and Table S4), ensuring exquisite accuracy during DNA replication. Below, we discuss the model that we propose for each step of the *C. crescentus* MutH-independent MMR process (Fig.8), which is based on *in vivo* assays characterizing the activity and the sub-cellular localization of wild-type and mutated MMR components described in this study and on models proposed in *Gammaproteobacteria* and *Bacilli* classes of bacteria (8,15).

**Figure 8:**
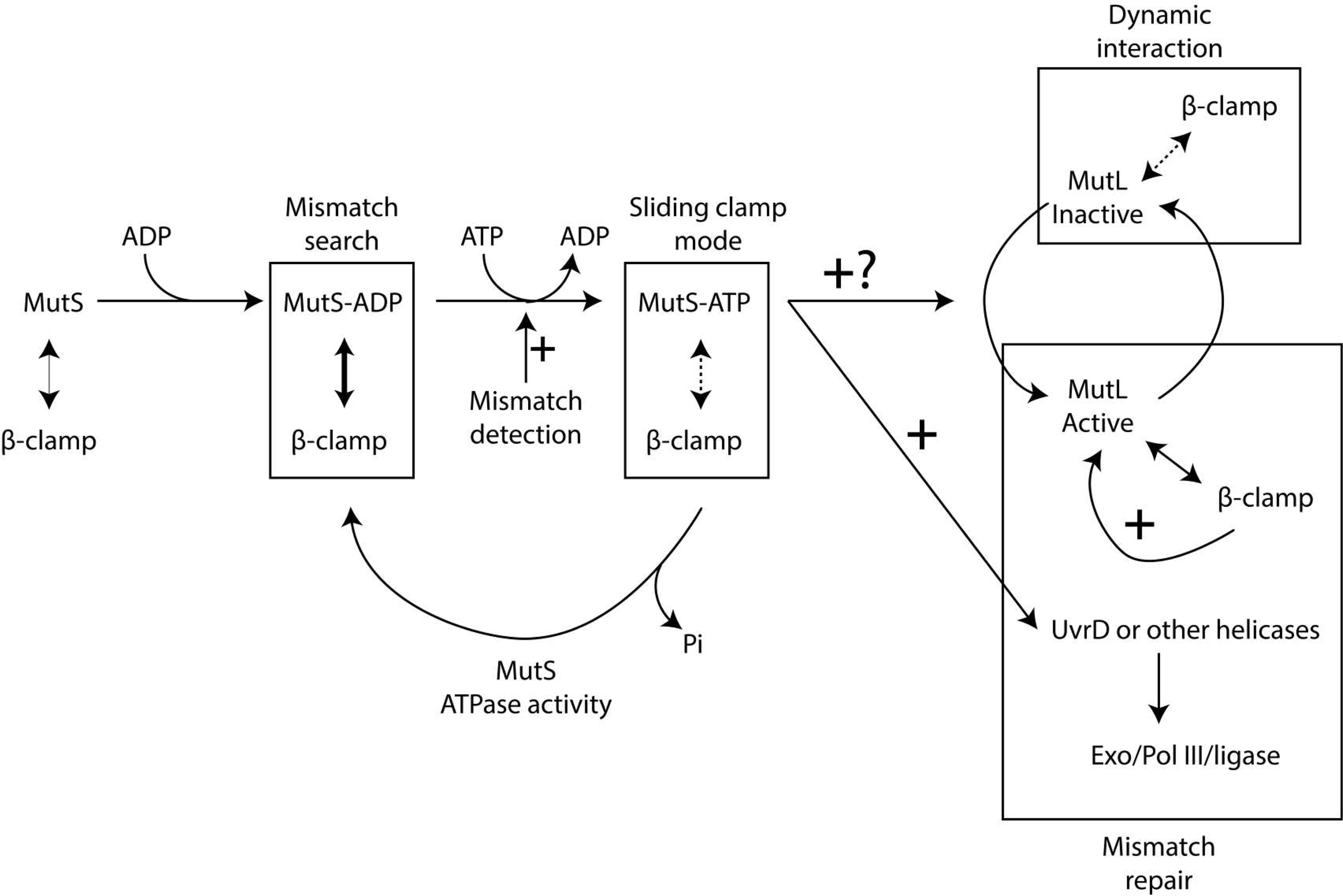
Model for the MMR process in *C. crescentus*. MutS-ADP binds to the β-clamp for 1D mismatch scanning during DNA replication. Mismatch detection by MutS triggers an ADP to ATP exchange and a conformational change in MutS, converting it into a sliding clamp that activates downstream MMR events. The ATP bound to MutS is then hydrolyzed, regenerating MutS-ADP that rapidly goes back to the replisome. MutL is dynamically recruited to the β-clamp during DNA replication and this interaction is needed for its activity as an endonuclease that nicks newly synthesized DNA strands. Mismatch detection by MutS most likely activates the latent endonuclease activity of MutL and/or helicases/exonucleases (Exo) needed for downstream events of the MMR process. The DNA polymerase III then resynthesizes the gap, while the ligase restores strand continuity.

### Mismatch searching by MutS in *C. crescentus*

Not surprisingly, our data shows that the *C. crescentus* MutS protein and its capacity to detect mis-paired bases through its conserved F_44_ motif plays a critical role in reducing the appearance of spontaneous mutations leading to antibiotic resistances (Fig.1 and Table S4). A major finding from this study is that fluorescently tagged but functional MutS appears to co-localize with the replisome during the whole S-phase of the *C. crescentus* cell cycle (Fig.3), in a manner that is dependent on a conserved 849DLPFL_853_ β-clamp binding motif found close to its C-terminal end (Fig.2), but independent of the frequency of mismatch occurrence (Fig.4). Compared with similar bulk fluorescence microscopy experiments done previously with fluorescently tagged MutS proteins from *E. coli* or *B. subtilis*, when it was found that only a minority (1-5%) of cells displayed clear fluorescent foci when no mutagen was added (36–38), our observations suggest that MutS is more stably associated with the β-clamp of the DNA polymerase in *C. crescentus* than it is in *B. subtilis* or *E. coli* cells. Interestingly, other conserved DnaN-interacting proteins, such as DnaE or HdaA, were also shown to bind to DnaN more efficiently in *C. crescentus* than in *E. coli* during recent *in vitro* experiments, suggesting that the *C. crescentus* β-clamp may display non-canonical properties (39). Furthermore, targeted mutagenesis of the conserved Walker A and B motifs of MutS indicate that ADP and ATP are important co-factors modulating the activity of MutS (Fig.1 and Table S4) and its capacity to interact with the replisome in *C. crescentus* (Fig.2). Overall, we propose a model in which MutS bound to ADP has the highest affinity for the replisome (Fig.2) to search for mismatches right behind the replication forks in a mostly 1D scanning mechanism during the whole S-phase of the cell cycle (Fig.8). Is this apparent 1D searching mode more efficient than a 3D searching mode? To address this important question, we isolated a mutant MutS(_849_AAAAA_853_) protein that was no more associated with the replisome *in vivo* (Fig.2) and found that it was almost as efficient in detecting and initiating the correction of mismatches than the wild-type protein (Fig.1 and Table S4). Thus, under standard growth conditions that do not promote replication errors, the spatial association of MutS with the replisome appears as strong but dispensible for MMR. Then, why is MutS associated with the replisome? One answer may be that this connection becomes important under non-standard growth conditions when the DNA polymerase may make more mistakes. Consistent with this proposal, we experienced severe difficulties when trying to bring a *dnaQ(G13E)* mutation into cells expressing *YFP-mutS(_849_AAAAA_853_)* as the only copy of *mutS* despite multiple attempts, generating only unstable and highly abnormal clones (data not shown).

### MMR activation upon mismatch detection in *C. crescentus*

According to the “sliding clamp” model for MMR (8,15), mismatch detection by MutS-ADP stimulates an ADP-to-ATP exchange, converting MutS into a “sliding clamp” with lower affinity for the replisome, which then quickly activates downstream MMR events. Consistent with this model, we found that a mutant *C. crescentus* MutS(E735A) protein, which is predicted to lack the ATPase activity, is significantly less often associated with the replisome (Fig.2), suggesting that MutS-ATP may form replisome-disconnected “sliding clamps” after mismatch detection in *C. crescentus* (Fig.8).

### Cleavage of newly synthesized DNA strands by MutL in *C. crescentus*

Unlike previous studies done on the *B. subtilis* MutL protein (38), we were lucky to be able to construct a fluorescently-tagged MutL protein that was almost fully functional in *C. crescentus* cells (Fig.1 and Table S4). Using this construct, we found that MutL is frequently, although not systematically, associated with the replisome (Fig.5), suggesting the existence of a dynamic mechanism recruiting and releasing MutL from the replisome during the S-phase of the cell cycle. Interestingly, this spatial association was shown to be dependent on a conserved _497_QTLLLP_502_ β-clamp binding motif located near the MutL C-terminus (Fig.6) (40), but independent of mismatch formation (Fig.4) or of the presence of a functional MutS protein (Fig.S7). Thus, contrarily to fluorescently-tagged MutL proteins previously analyzed in *B. subtilis* cells (37), the *C. crescentus* MutL protein appears to be regularly recruited to the replisome by the β-clamp even if MutS does not detect mismatches. Since we found that an inactive MutL(D472N) protein lacking the endonuclease activity (Fig.1 and Table S4) is significantly stabilized at the replisome compared to the wild-type protein (Fig.5A), we propose that MutL cuts the newly synthesized DNA strand when it is located at the replisome (Fig.8). Consistent with this proposal, we found that a MutL(_497_ATLAAP_502_) protein that is no more recruited to the replisome (Fig.5A) is totally inactive (Fig.1 and Table S4) *in vivo*. Although we cannot rule out the possibility that the DnaN-induced *C. crescentus* MutL protein may cut the newly synthesized DNA strand regularly independently of mismatch detection by MutS, as recently suggested by some *in vitro* assays using the *B. subtilis* MutL protein (19), we favour a model in which MutS-ATP triggers this cleavage by a MutL-DnaN complex at the replisome (Fig.8). How MutL recognizes the newly synthesized DNA strand that needs to be repaired remains unknown in all organisms lacking MutH/Dam. Although a vast majority of *Alphaproteobacteria* possess an orphan CcrM DNA methyltransferase that methylates adenines in 5’GANTC3’ motifs, we showed years ago that it does not play a role similar to Dam in *Gammaproteobacteria*, as a *C. crescentus* mutant lacking *ccrM* is not a mutator strain (41,42). Instead, it is tempting to speculate that the spatial association between MutL and the replisome may contribute to strand discrimination during the *C. crescentus* MMR process.

### Downstream steps of the MMR process in *C. crescentus*

Once the mismatch-containing strand is cut by MutL in bacteria lacking MutH/Dam, it is unclear which helicase is responsible for strand separation prior to digestion by exonucleases (15). Since *C. crescentus* has a protein homologous to UvrD (43), which plays an important role at that step during the *E.coli* MutH-dependent MMR process, we tested whether it may play a similar role in *C. crescentus*. Although *uvrD* mutants are mutator strains (Fig.1, Table S4 and (30)), our data suggests that UvrD is not the helicase involved in *C. crescentus* MMR (Fig.7), or that there exist more than one helicase involved with significant functional redundancy (Fig.8). Clearly, understanding how late steps of the MMR process take place in a variety of different bacteria is an interesting avenue for future research and may again contribute to understanding why MMR is spatially associated with DNA replication in so many organisms.

## SUPPLEMENTARY DATA

Supplementary information is available.

## AKNOWLEDGEMENTS

We would like to thank past and current members of the Collier team for helpful discussions, Noémie Matthey for useful comments on the manuscript and Renske van Raaphorst for some help to use the Oufti software.

