## Supplementary material for "Spatial coupling between DNA replication and mismatch repair in *Caulobacter crescentus*"

**Content:**

**Supplementary Tables** (page 2)

Table S1 (page 2)

Table S2 (page 3)

Table S3 (page 4)

Table S4 (page 5)

**Supplementary Figures with legends** (page 6)

Fig.S1 (page 6)

Fig.S2 (page 8)

Fig.S3 (page 9)

Fig.S4 (page 10)

Fig.S5 (page 11)

Fig.S6 (page 12)

Fig.S7 (page 13)

Fig.S8 (page 14)

**Supplementary Materials and Methods** (page 15)

**Supplementary References** (page 16)

### Supplementary Tables

**Table S1: Oligonucleotides used in this study.** Restriction enzyme recognition sites used for cloning purposes are underlined.

| Name | Sequence (5'→3') | Use |
|---|---|---|
| **CT1** | GGTAAGCTTGGTCCGCGTGGGGATCTCTAC (HindIII) | Construction of pNPTS138-ΔmutS |
| **CT2** | CCCGGATCCGGGCGGCGTTAAGGAATTACAC (BamHI) | |
| **CT3** | CCCGGATCCTAGCGGGTACCCATATAGA (BamHI) | |
| **CT4** | CGGGCTAGCAACCCAGATCCCACAACGCATAC (NheI) | |
| **CT8** | CGGGCTAGCCGAGACGATGGATGACAAACACC (NheI) | Construction of pNPTS138-ΔmutL |
| **CT10** | GGTAAGCTTATGCCCATCCGCCGCCTGCCG (HindIII) | |
| **CT11** | CCCGGATCCCATGGCCTGGCGCTTGATCTCC (BamHI) | |
| **CT12** | CCCGGATCCTGAGGCCAGAGCGCCCCCTAC (BamHI) | |
| **TC46** | GTCGGATCCGCCTGATCGACAGGGCGAACCA (BamHI) | Construction of pNPTS138-ΔuvrD |
| **TC47** | CGGGAATTCCGCCGAAAAGCGAGCGTTCAGC (EcoRI) | |
| **TC63** | ATAAAGCTTATGTTTCCCGACACCGACTCGC (HindIII) | |
| **TC64** | CGTGGATCCGCGGTTCTTCCAGCCGTCGATG (BamHI) | |
| **TC18** | GCGAAGCTTGGTCCGCGTGGGGATCTCTA (HindIII) | Construction of pNPTS138-PmutS-YFP-MutS |
| **TC19** | GTGGGATCCCATGACGCCCGGAAACTACAG (BamHI) | |
| **TC20** | GAAGGATCCATGGTGAGCAAGGGCGAGGAG (BamHI) | |
| **TC21** | CTCGAATTCGGACAGCTCGACCGACGCCA (EcoRI) | |
| **TC42** | TGACTGACCGCGGCGGCGGCGGCGGCGTCCAGCTTGG | Construction of pNPTS138- MutS($_{849}$AAAAA$_{853}$) and pXYFPN4- MutS($_{849}$AAAAA$_{853}$) |
| **TC43** | CCAAGCTGGACGCCGCCGCCGCCGCCGCGGTCAGTCA | |
| **TC52** | AACAAGCTTGGTAAGTCGACCTTCCTGCGCC, HindIII | Construction of pNPTS138- MutS($_{849}$AAAAA$_{853}$) |
| **TC53** | GCGGAATTCGCTTGTAGGGCCGCAGGAACAG, EcoRI | |
| **TC34** | CGAAAGCTTCTTTCCCGGCGTCGTTGTGGA (HindIII) | Construction of pNPTS138-DnaQ(G13E) |
| **TC35** | TTTGGGTCAAACTCGGTGGTTTCG | |
| **TC36** | CGAAACCACCGAGTTTGACCCAAA | |
| **TC37** | GTGGGATCCGCCATGGCCCCTGTCGAATGG (BamHI) | |
| **TC14** | GGAAAGCTTCCAGCAGGCTCAGAATCCGAA (HindIII) | Construction of pNPTS138-PmutL-YFP-MutL |
| **TC15** | GCGGGATCCCATCAAGCGGACTTTCACGGG (BamHI) | |
| **TC16** | GACGGATCCATGGTGAGCAAGGGCGAGGA (BamHI) | |
| **TC17** | GGCGAATTCGGACTTCATGAATTTCAGGCG (EcoRI) | |
| **JC155** | CGGGTACCATGAACGCCCACGCCACGCCGACC (KpnI) | Construction of pXYFPN4-MutS, pXYFPN4-MutS($_{849}$AAAAA$_{853}$), pXYFPN4-MutS(F44A), pXYFPN4-MutS(K661M), pXYFPN4-MutS(E735A) |
| **JC156** | GGCTAGCCTAGGCCGTGAGCAGACCCTTAAGG (NheI) | |
| **TC27** | CAGCTCGTAGGCATCGCCCATGC | Construction of pXYFPN4-MutS(F44A) |
| **TC28** | GCATGGGCGATGCCTACGAGCTG | |
| **TC23** | GAAGGTCGACATACCGGCCATG | Construction of pXYFPN4-MutS(K661M) |
| **TC24** | CATGGCCGGTATGTCGACCTTC | |
| **TC38** | ATCCTGGACGCCATCGGCCGGG | Construction of pXYFPN4-MutS(E735A) |
| **TC39** | CCCGGCCGATGGCGTCCAGGAT | |
| **CoC-7_1** | TCGGTACCATGCCCATCCGCCGCCTGCCG (KpnI) | Construction of pXYFPN4-MutL, pXYFPN4-MutL($_{497}$ATLAAP$_{502}$), pXYFPN4-MutL(D472N) and pXYFPN4-MutL(D472N, $_{497}$ATLAAP$_{502}$) |
| **CoC-8_4** | GGAGCTCTGGCCTCACCGCCGCCCGAACAGC (SacI) | |
| **TC40** | ACCTCGGGGGCGGCCAGGGTGGCGCGGGTC | Construction of pXYFPN4- MutL($_{497}$ATLAAP$_{502}$) and pXYFPN4-MutL(D472N, $_{497}$ATLAAP$_{502}$) |
| **TC41** | GACCCGCGCCACCCTGGCCGCCCCCGAGGT | |
| **TC31** | CGCGTGCTGGTTGACAATGAC | Construction of pXYFPN4-MutL(D472N) and pXYFPN4-MutL(D472N, $_{497}$ATLAAP$_{502}$) |
| **TC32** | GGTCATTGTCAACCAGCACGCG | |
| **TC54** | TAGGTACCATGTTTCCCGACACCGACTCGC (KpnI) | Construction of pXYFPN4-UvrD |
| **TC55** | CCCGCTAGCTCAGGCCTTCTCGACGAAGCTG (NheI) | |

### Table S2: Plasmids used in this study.

| Plasmid name | Description | Source |
|---|---|---|
| pNPTS138 | Suicide vector carrying the *sacB* gene | D. Alley, unpublished |
| pNPTS138-ΔmutS | pNPTS138 derivative used to create the *ΔmutS* deletion strain | This study |
| pNPTS138-ΔmutL | pNPTS138 derivative used to create the *ΔmutL* deletion strain | This study |
| pNPTS138-ΔuvrD | pNPTS138 derivative used to create the *ΔuvrD* deletion strain | This study |
| pNPTS138-PmutS-YFP-MutS | pNPTS138 derivative used to replace *mutS* by *yfp-mutS* (under the control of native *mutS* promoter) | This study |
| pNPTS138-DnaQ(G13E) | To replace native *dnaQ (CCNA_00005)* with *dnaQ(G13E)* allele | This study |
| pNPTS138-MutS($_{849}$AAAAA$_{853}$) | To replace native *mutS* with the *mutS*($_{849}$AAAAA$_{853}$) allele | This study |
| pNPTS138-PmutL-YFP-MutL | pNPTS138 derivative used to replace *mutL* by *yfp-mutL* (under the control of native *mutL* promoter) | This study |
| pXYFPN4 | For integrating constructs encoding N-terminal YFP fusions under the control of the native *xylX* promoter | (1) |
| pXYFPN4-MutS | *mutS* cloned into pXYFPN4 | This study |
| pXYFPN4-MutS($_{849}$AAAAA$_{853}$) | *mutS*($_{849}$AAAAA$_{853}$) cloned into pXYFPN4 | This study |
| pXYFPN4-MutS(F44A) | *mutS(F44A)* cloned into pXYFPN4 | This study |
| pXYFPN4-MutS(K661M) | *mutS(K661M)* cloned into pXYFPN4 | This study |
| pXYFPN4-MutS(E735A) | *mutS(E735A)* cloned into pXYFPN4 | This study |
| pXYFPN4-MutL | *mutL* cloned into pXYFPN4 | This study |
| pXYFPN4-MutL($_{497}$ATLAAP$_{502}$) | *mutL*($_{497}$ATLAAP$_{502}$) cloned into pXYFPN4 | This study |
| pXYFPN4-MutL(D472N) | *mutL(D472N)* cloned into pXYFPN4 | This study |
| pXYFPN4-MutL(D472N, $_{497}$ATLAAP$_{502}$) | *mutL*(D472N, $_{497}$ATLAAP$_{502}$) cloned into pXYFPN4 | This study |
| pXYFPN4-UvrD | *uvrD* cloned into pXYFPN4 | This study |

### Table S3: Strains used in this study.

| **Strain name** | **Genotype** | **Source** |
|---|---|---|
| *Escherichia coli* | | |
| TOP10 | *F- mcrA Δ(mrr-hsdRMS-mcrBC) φ80lacZΔM15 ΔlacX74 nupG recA1 araD139 Δ(ara-leu)7697 galE15 galK16 rpsL($Str^R$) endA1 $λ^-$* | Invitrogen (USA) |
| *Caulobacter crescentus* | | |
| NA1000 | Synchronizable derivative of wild-type strain CB15 (CB15N) | (2) |
| JC1427 | NA1000 *ΔmutS* | This study |
| JC1426 | NA1000 *ΔmutL* | This study |
| JC1575 | NA1000 *ΔmutS ΔmutL* | This study |
| JC1847 | NA1000 *ΔuvrD* | This study |
| JC1986 | NA1000 *ΔuvrD ΔmutL* | This study |
| JC1784 | NA1000 *YFP-mutS* | This study |
| JC1433 | NA1000 Δ*mutS Pxyl::YFP-mutS* | This study |
| JC577 | NA1000 *dnaN-CFP* | (3) |
| JC1625 | NA1000 *dnaN-CFP* Δ*mutS Pxyl::YFP-mutS* | This study |
| JC1770 | NA1000 Δ*mutS Pxyl::YFP-mutS*($_{849}$AAAAA$_{503}$) | This study |
| JC1799 | NA1000 *mutS*($_{849}$AAAAA$_{503}$) | This study |
| JC1678 | NA1000 *dnaQ(G13E)* | This study |
| JC1724 | NA1000 Δ*mutS Pxyl::YFP-mutS* pNPTS138-DnaQ(G13E) integrated at *dnaQ* locus | This study |
| JC1666 | NA1000 Δ*mutS Pxyl::YFP-mutS(F44A)* | This study |
| JC1669 | NA1000 *dnaN-CFP* Δ*mutS Pxyl::YFP-mutS(F44A)* | This study |
| JC1665 | NA1000 Δ*mutS Pxyl::YFP-mutS(K661M)* | This study |
| JC1739 | NA1000 Δ*mutS Pxyl::YFP-mutS(E735A)* | This study |
| JC1668 | NA1000 *dnaN-CFP* Δ*mutS Pxyl::YFP-mutS(K661M)* | This study |
| JC1740 | NA1000 *dnaN-CFP* Δ*mutS Pxyl::YFP-mutS(E735A)* | This study |
| JC1769 | NA1000 *YFP-mutL* | This study |
| JC1825 | NA1000 Δ*mutL Pxyl::YFP-mutL* | This study |
| JC1812 | NA1000 *dnaN-CFP* Δ*mutL Pxyl::YFP-mutL* | This study |
| JC1805 | NA1000 *dnaN-CFP* Δ*mutS* Δ*mutL Pxyl::YFP-mutL* | This study |
| JC1749 | NA1000 Δ*mutL Pxyl::YFP-mutL*($_{497}$ATLAAP$_{502}$) | This study |
| JC1750 | NA1000 *dnaN-CFP* Δ*mutL Pxyl::YFP-mutL*($_{497}$ATLAAP$_{502}$) | This study |
| JC1667 | NA1000 Δ*mutL Pxy::YFP-mutL(D472N)* | This study |
| JC1670 | NA1000 *dnaN-CFP* Δ*mutL Pxy::YFP-mutL(D472N)* | This study |
| JC1894 | NA1000 *dnaN-CFP* Δ*mutS* Δ*mutL Pxy::YFP-mutL(D472N)* | This study |
| JC1753 | NA1000 *dnaN-CFP* Δ*mutL Pxyl::YFP-mutL*(D472N, $_{497}$ATLAAP$_{502}$) | This study |
| JC1870 | NA1000 Δ*uvrD Pxyl::YFP-uvrD* | This study |
| JC1846 | NA1000 *Pxyl::YFP-uvrD* | This study |
| JC2211 | NA1000 *Pxyl::YFP-uvrD* pNPTS138-DnaQ(G13E) integrated at *dnaQ* locus | This study |
| JC1977 | NA1000 Δ*mutS Pxyl::YFP-uvrD* | This study |

**Table S4: Comparison of spontaneous mutation rates of *C. crescentus* strains used in this study.** The symbols * and # indicate that the mutation rate is statistically significantly different or equivalent, respectively, to that of the NA1000 (WT) reference strain (for Zones A and D), of the JC1433 (Δ*mutS Pxyl::YFP-mutS)* strain (Zone B) or of the JC1825 *(ΔmutL Pxyl::YFP-mutL)* strain (Zone C). Student's *t*-tests were used to estimate statistical significance: differences were considered as significant (*) if $P<0.05$. The "relative mutation rate" value, facilitating comparisons, was calculated setting the spontaneous mutation rate of the NA1000 (WT) strain to a value of 1 (arbitrary units).

| Zone | Strain number | Genotype (disturbed activity) | Number of Rif$^R$ colonies * 10$^{-9}$ total colonies [±standard deviation] | Relative mutation rate [% of DNA repair activity] |
|---|---|---|---|---|
| A | NA1000 | WT | 8.3 [±2.8] | 1 [=100%] |
| | JC1427 | *ΔmutS* | 998.3 [±212.9]* | 120.3 [=0%] |
| | JC1426 | *ΔmutL* | 916.7 [±312.4]* | 110.4 [=0%] |
| | JC1575 | *ΔmutS ΔmutL* | 1152.7 [±423.1]* | 138.9 |
| | JC1847 | *ΔuvrD* | 540.1 [±95.0]* | 65.1 [=0%] |
| | JC1986 | *ΔuvrD ΔmutL* | 1979.3 [±460.0]* | 238.5 |
| | JC577 | *dnaN-CFP* | 55.8 [±33.6]# | 6.7 |
| | JC1784 | *YFP-mutS* | 52.8 [±21.4]# | 6.4 [95.5%] |
| | JC1799 | *mutS*($_{849}$*AAAAA*$_{853}$) (β-clamp binding) | 56.8 [±24.3]* | 6.8 [95.1%] |
| | JC1678 | *dnaQ(G13E)* | 3973.3 [±1466.0]* | 478.7 |
| | JC1769 | *YFP-mutL* | 148.6 [±78.1]# | 17.9 [84.6%] |
| | JC2212 | *mutL*($_{497}$*ATLAAP*$_{502}$) (β-clamp binding) | 932.7 [±315.6]* | 112.4 [~0%] |
| B | JC1433 | Δ*mutS Pxyl::YFP-mutS* | 24.2 [±9.9] | 2.9 [98.4%] |
| | JC1770 | Δ*mutS Pxyl::YFP-mutS*($_{849}$*AAAAA*$_{853}$) (β-clamp binding) | 45.0 [±21.2]# | 5.4 [96.3%] |
| | JC1666 | Δ*mutS Pxyl::YFP-mutS(F44A)* (mismatch recognition) | 843.0 [±313.2]* | 101.6 [15.7%] |
| | JC1665 | Δ*mutS Pxyl::YFP-mutS(K661M)* (nucleotide binding and ATP hydrolysis) | 792.3 [±75.0]* | 95.5 [20.8%] |
| | JC1739 | Δ*mutS Pxyl::YFP-mutS(E735A)* (ATP hydrolysis) | 917.0 [±340.0]* | 110.5 [8.2%] |
| C | JC1825 | Δ*mutL Pxyl::YFP-mutL* | 56.3 [±28.5] | 6.8 [94.7%] |
| | JC1749 | Δ*mutL Pxyl::YFP-mutL*($_{497}$*ATLAAP*$_{502}$) (β-clamp binding) | 751.8 [±71.4]* | 90.6 [17.2%] |
| | JC1667 | Δ*mutL Pxyl::YFP-mutL(D472N)* (endonuclease) | 904.3 [±204.2]* | 109.0 [0.4%] |
| D | JC1870 | Δ*uvrD Pxyl::YFP-uvrD* | 102.5 [±39.5]# | 12.3 [82.3%] |

### Supplementary Figures:

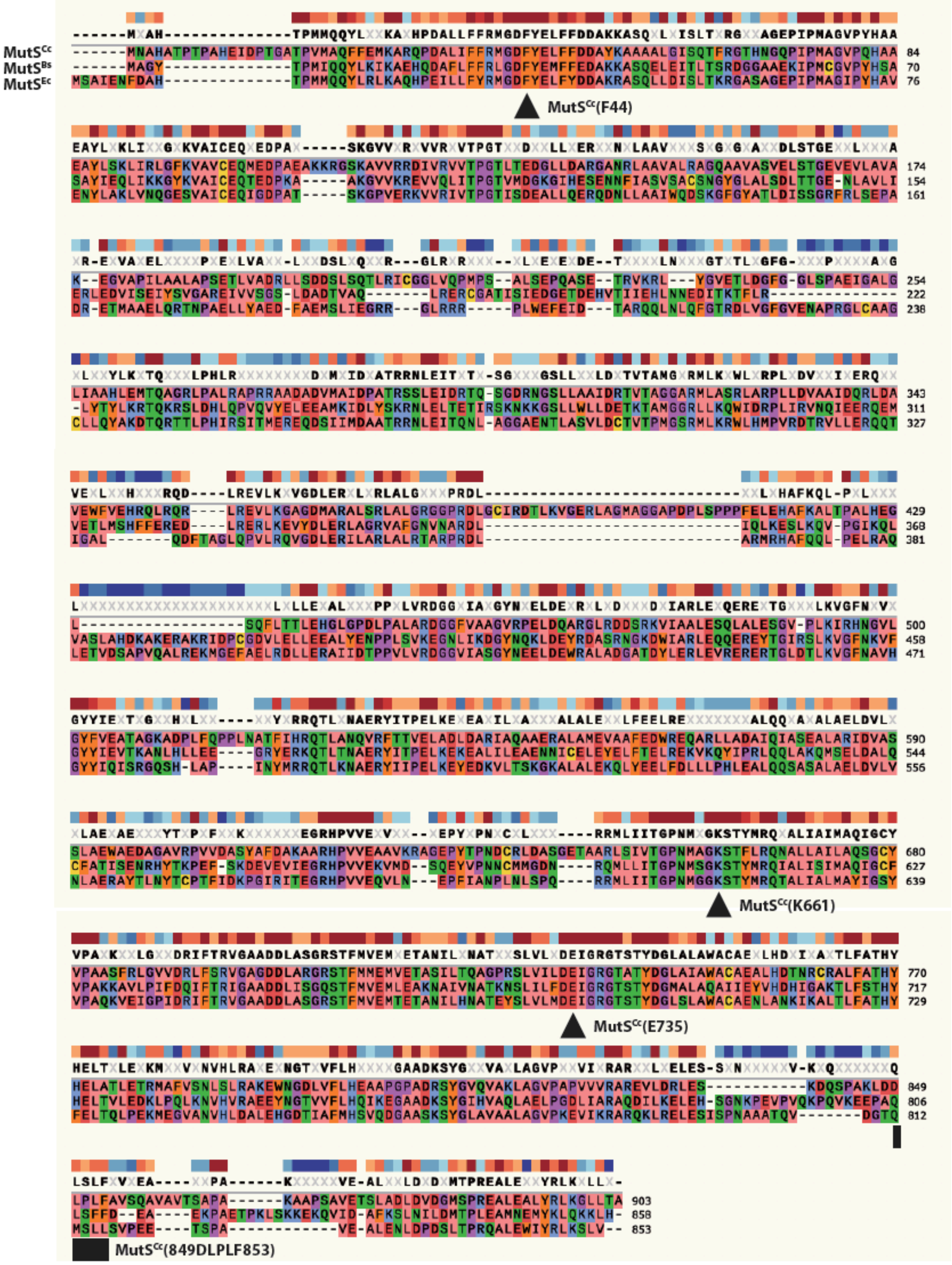


**Figure S1: Similarities between selected MutS proteins.** Amino-acid sequences of MutS homologs were downloaded from NCBI for *Caulobacter crescentus* NA1000 strain (MutS$^{Cc}$), *Bacillus subtilis* PY79 strain (MutS$^{Bs}$) and *Escherichia coli* MG1655 strain (MutS$^{Ec}$).

Alignments were done using the SnapGene software (from Insightful Science; available at snapgene.com). MutS$^{Cc}$ residues/motifs of particular interest are indicated under the three sequences. The F44 residue belongs to a $_{42}\underline{G}D\underline{FY}ELFFD\underline{DA}_{52}$ motif where underlined residues are part of the consensus motif for mismatch recognition by MutS proteins (4). The K661 residue belongs to a $_{655}$GPNMAGKS$_{662}$ Walker A motif supposedly involved in ATP/ADP binding and whose consensus is GxxxxGK[T/S] (x can be any amino acid) (5). The E735 residue belongs to a $_{689}$LVLMDE$_{694}$ Walker B motif supposedly involved in ATP hydrolysis and whose consensus is hhhhDE (h is a hydrophobic amino acid) (5). The $_{849}$DLPFL$_{853}$ motif shows similarities with the β-clamp binding motifs MutS$^{Bs}$ ($_{806}$QLSFF$_{810}$) (6)and MutS$^{Ec}$ ($_{812}$QMSLL$_{816}$) (7).

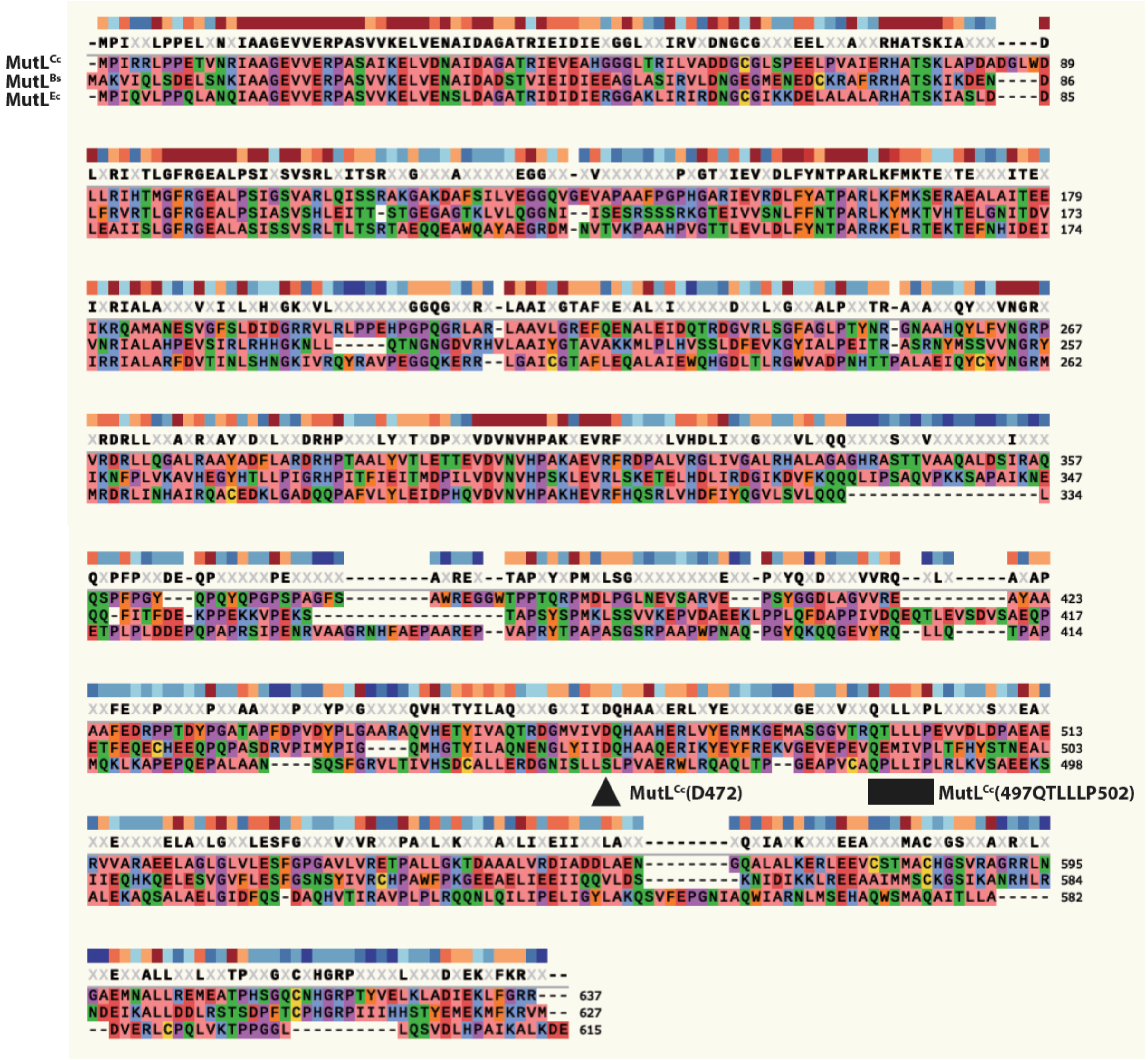


**Figure S2: Similarities between selected MutL proteins.** Amino-acid sequences of MutL homologs were downloaded from NCBI for *Caulobacter crescentus* NA1000 strain (MutS$^{Cc}$), *Bacillus subtilis* PY79 strain (MutS$^{Bs}$) and *Escherichia coli* MG1655 strain (MutS$^{Ec}$). Alignments were done using the SnapGene software. MutL$^{Cc}$ residues/motifs of particular interest are indicated under the three sequences. The D472 residue belongs to the predicted ${}_{472}\underline{\mathrm{DQHA}}\mathrm{AH}\underline{\mathrm{E}}\mathrm{RLVY}\underline{\mathrm{E}}_{483}$ endonuclease domain (underlined letters indicate conserved residues according to (8)). The ${}_{497}$QTLLLP${}_{502}$ motif shows similarities with the previously proposed Qxh(L/I)xP consensus β-clamp binding motif for MutL proteins (9).

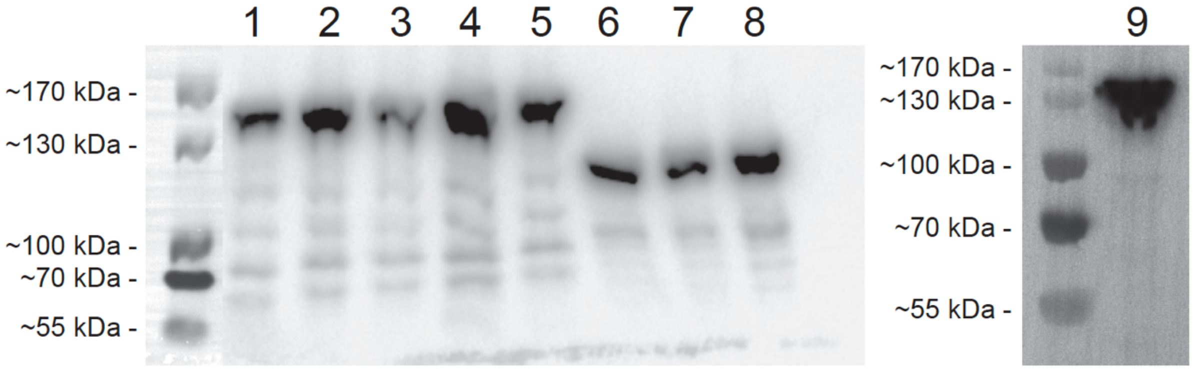


**Figure S3: Immunoblot detection of YFP-tagged MutS, MutL and UvrD derivatives from *C. crescentus* cell extracts.** Cells from strains JC1433 (Δ*mutS Pxyl::YFP-mutS*) (Lane 1), JC1665 (Δ*mutS Pxyl::YFP-mutS(K661M)*) (Lane 2), JC1666 (Δ*mutS Pxyl::YFP-mutS(F44A)*) (Lane 3), JC1739 (Δ*mutS Pxyl::YFP-mutS(E735A)*) (Lane 4), JC1770 (Δ*mutS Pxyl::YFP-mutS(*$_{849}$*AAAAA*$_{853}$*)*) (Lane 5), JC1825 (Δ*mutL Pxyl::YFP-mutL*) (Lane 6), JC1667 (Δ*mutL Pxyl::YFP-mutL(D472N)*) (Lane 7), JC1749 (Δ*mutL Pxyl::YFP-mutL(*$_{497}$*ATLAAP*$_{502}$*)*) (Lane 8) and JC1870 (Δ*uvrD Pxyl::YFP-uvrD*) (Lane 9) were cultivated overnight in PYE medium before the culture was diluted into M2G medium. Once the $OD_{660nm}$ reached ~0.3 (mid-exponential phase), 0.3% xylose was added. Cell extracts were then prepared when cultures reached an $OD_{660nm}$ of ~0.5 (exponential phase) and proteins were separated by SDS-PAGE. YFP-tagged proteins were then detected by immunoblotting using anti-GFP antibodies. MW corresponds to the molecular weight in kDa. This figure shows that the YFP moiety of each fusion protein is not cleaved off.

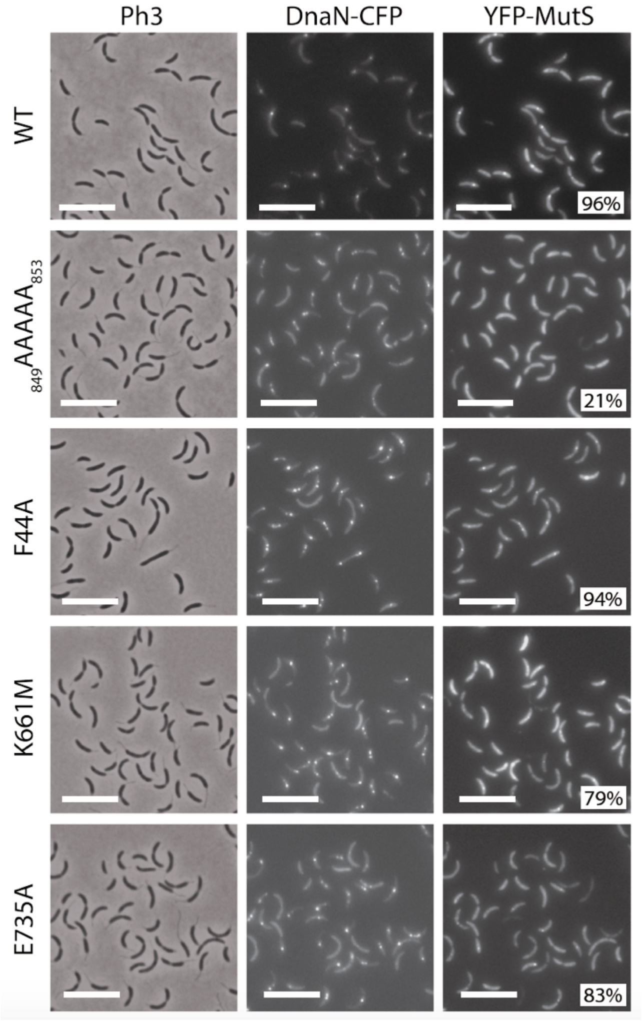


**Figure S4: YFP-MutS foci co-localize with the replisome in *C. crescentus*.** Subcellular localization of DnaN-CFP and derivatives of YFP-MutS in Δ*mutS* cells. Strains JC1625 (Δ*mutS dnaN-CFP Pxyl::YFP-mutS*), JC1771 (Δ*mutS dnaN-CFP Pxyl::YFP-mutS*($_{849}$*AAAAA*$_{853}$)), JC1669 (Δ*mutS dnaN-CFP Pxyl::YFP-mutS(F44A)*), JC1668 (Δ*mutS dnaN-CFP Pxyl::YFP-mutS(K661M)*) and JC1740 (Δ*mutS dnaN-CFP Pxyl::YFP-mutS(E735A)*) were cultivated into PYE medium and then transferred into M2G medium. 0.3% xylose was added to cultures when they reached an $OD_{660nm}$~0.3. Cells were then imaged by fluorescence microscopy when the $OD_{660nm}$ reached ~0.5. The % indicated onto images corresponds to the average proportion of distinct MutS-YFP foci (intensity >2-fold above background) that are co-localized with DnaN-CFP foci (using values obtained from three independent experiments). The white scale bar corresponds to 8μm.

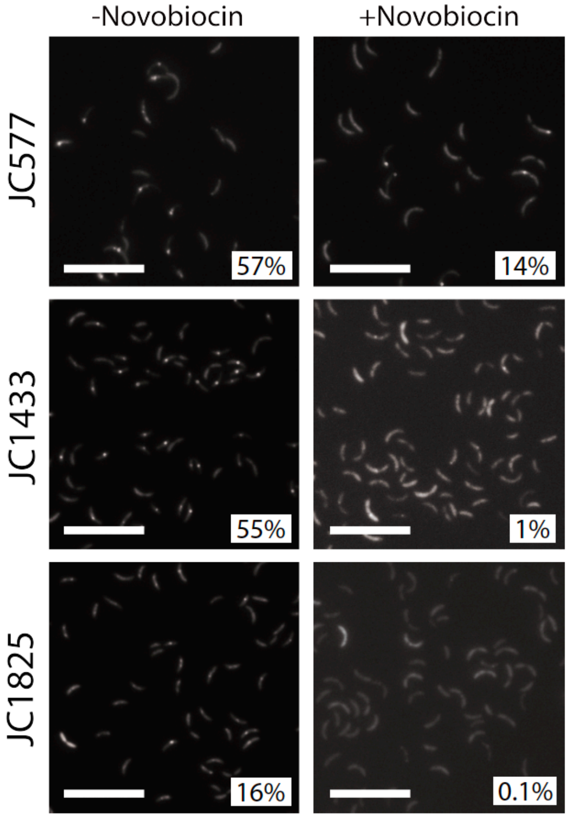


**Figure S5: Replication inhibition following novobiocin treatment largely dissociates DnaN-CFP, YFP-MutS and YFP-MutL foci.** Strains JC577 (*dnaN-CFP*), JC1433 (Δ*mutS Pxyl::YFP-mutS*) and JC1825 (Δ*mutL Pxyl::YFP-mutL*) were cultivated in PYE medium and then transferred into M2G medium. When the $OD_{660nm}$ reached ~0.3, cultures were split into two and 100 µg/mL of novobiocin was added into one of the sub-cultures. 0.3% xylose was added at the same time into each sub-cultures of strains JC1433 and JC1825. Once the $OD_{660nm}$ of the sub-culture without novobiocin reached 0.5, cells were imaged by fluorescence microscopy using CFP (for DnaN-CFP) or YFP (for YFP-MutS or YFP-MutL) illumination. Representative images are shown here. % indicated under each image correspond to the average proportion of cells displaying CFP or YFP foci (intensity >2-fold above background) as counted during three independent experiments. The white scale bar corresponds to 11µm.

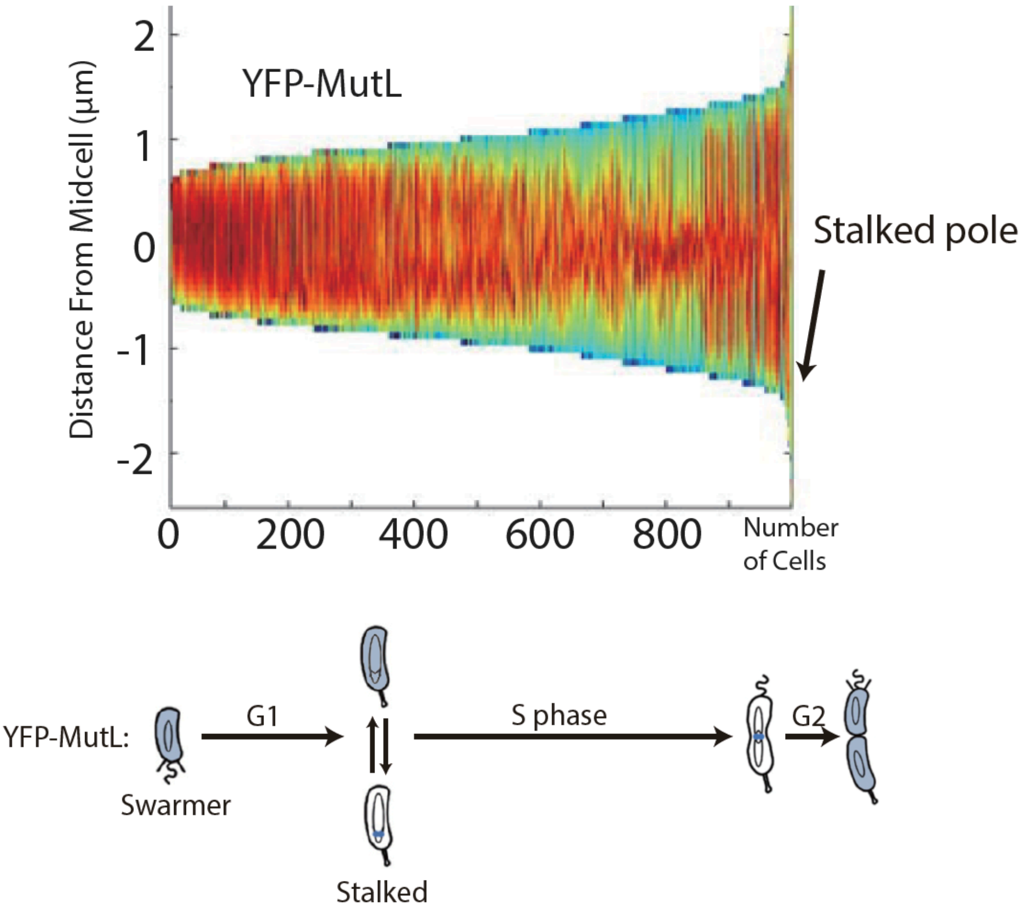


**Figure S6: YFP-MutL is enriched at mid-cell towards the end of the S-phase of the *C. crescentus* cell cycle.** Demograph showing the subcellular localization of YFP-MutL in Δ*mutL* cells sorted as a function of their size. JC1825 (Δ*mutL Pxyl::YFP-mutL*) cells were cultivated and imaged as described for Fig.5A. Short cells correspond to G1/swarmer cells, while intermediate and longer cells correspond to stalked and pre-divisional S-phase cells, respectively. The schematic under the demograph shows the *C. crescentus* cell cycle and the blue color highlights where YFP-MutL seems to be localized as a function of the cell cycle.

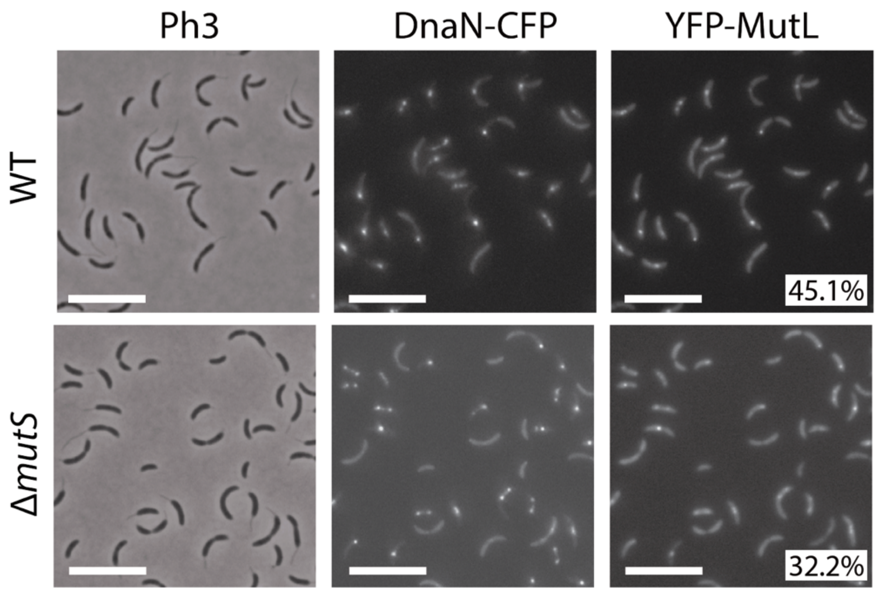


**Figure S7: YFP-MutL is recruited to the replisome independently of MutS.** Subcellular localization of DnaN-CFP and YFP-MutL in Δ*mutL* cells with or without *mutS*. Strains JC1812 (*dnaN-CFP* Δ*mutL Pxyl::YFP-mutL*) labelled "WT" and JC1805 (*ΔmutS* Δ*mutL dnaN-CFP Pxyl::YFP-mutL*) labelled *"ΔmutS*" were cultivated into PYE medium and then transferred into M2G medium. 0.3% xylose was added to cultures when they reached an $OD_{660nm}$~0.3. Cells were then imaged by fluorescence microscopy when the $OD_{660nm}$ reached ~0.5. The % indicated onto images corresponds to the average proportion of cells with distinct MutL-YFP foci (intensity >2-fold above average background) using values obtained from three independent experiments. The white scale bar corresponds to 8µm.

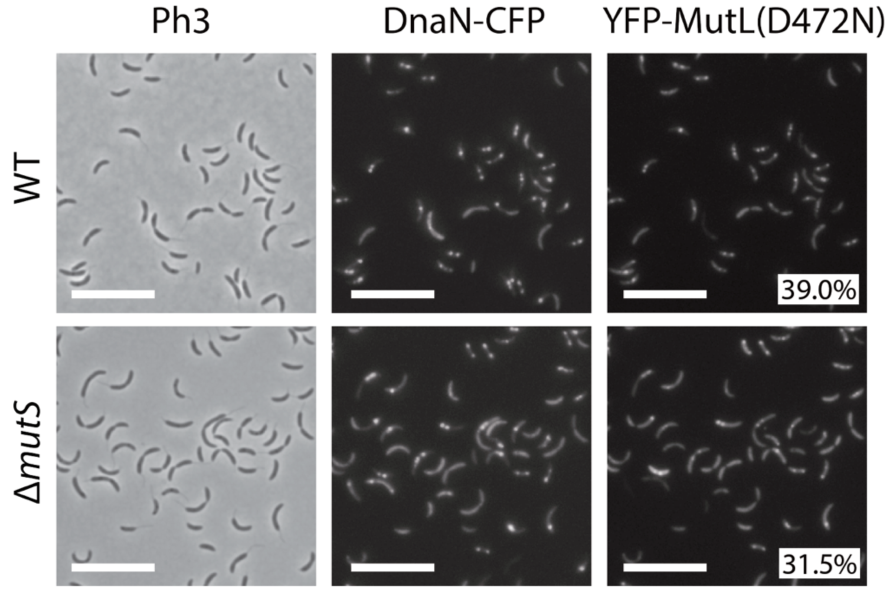


**Figure S8: YFP-MutL(D472N) is stabilized at the replisome even in the absence of MutS.** Subcellular localization of DnaN-CFP and YFP-MutL(D472N) in Δ*mutL* cells with or without *mutS*. Strains JC1670 (Δ*mutL dnaN-CFP Pxyl::YFP-mutL (D472N)*) and JC1894 (*ΔmutS ΔmutL dnaN-CFP Pxyl::YFP-mutL (D472N)*) were cultivated into PYE medium and then transferred into M2G medium. 0.3% xylose was added to cultures when they reached an $OD_{660nm}$~0.3. Cells were then imaged by fluorescence microscopy when the $OD_{660nm}$ reached ~0.5. The % indicated onto images corresponds to the average proportion of cells with distinct MutL-YFP(D472N) foci (intensity >2-fold above average background) using values obtained from three independent experiments. The white scale bar corresponds to 11µm.

### Supplementary Material and Methods

#### Plasmids constructions

*Derivatives of pNPTS138:*

To construct pNPTS138-ΔmutS, pNPTS138-ΔmutL and pNPTS138-ΔuvrD, ~500 bp sequences upstream and downstream of the corresponding ORF were amplified by PCR from wild-type *C. crescentus* chromosomal DNA using the primer pairs listed in Table S1. The two fragments were subsequently digested by restriction enzymes (as indicated in Table S1) and cloned into the pNPTS138 suicide vector between corresponding restriction sites by triple ligation.

To construct pNPTS138-DnaQ(G13E), pNPTS138-MutS($_{849}$AAAAA$_{853}$) and pNPTS138-MutL($_{497}$ATLAAP$_{503}$), overlap extension PCR were used to construct ~1 kb gene fragment with the required mutations in *dnaQ*, *mutS* or *mutL*. The PCR consisted of two consecutive PCR reactions, with four primers (listed in Table S1).

To construct pNPTS138-PmutS-YFP-mutS and pNPTS138-PmutL-YFP-mutL, two primer pairs (listed in Table S1) were used to amplify the ~500 bp promoter regions (PmutS and PmutL, respectively) and the protein fusion regions (YFP-mutS and YFP-mutL, respectively). The two fragments were subsequently digested by the indicated restriction enzymes (Table S1) and cloned into the pNPTS138 suicide vector between corresponding restriction sites by triple ligation.

*Derivatives of pXYFPN4:*

To construct pXYFPN4-MutS, pXYFPN4-MutL and pXYFPN4-UvrD, the corresponding ORFs were amplified by PCR from wild-type *C. crescentus* chromosomal DNA using the primer pairs listed in Table S1, digested by the indicated restriction enzymes and cloned into pXYFPN4.

To construct pXYFPN4-MutS(F44A), pXYFPN4-MutS(K661M), pXYFPN4-MutS(E735A), pXYFPN4-MutS($_{849}$AAAAA$_{853}$), pXYFPN4-MutL(D472N), pXYFPN4-MutL($_{497}$ATLAAP$_{502}$) and pXYFPN4-MutL(D472N , $_{497}$ATLAAP$_{502}$) overlap PCR was used to construct a point mutation in the *mutS* or *mutL* genes. Two consecutive PCR reactions were performed using four primers (listed in Table S1). First, two DNA fragments were amplified from plasmid pXYFPN4-mutS (for *mutS* alleles) or pXYFPN4-mutL (for *mutL* alleles). Then, equimolar amounts of both products were mixed and amplified. The resulting fragments were digested by restriction enzymes (as indicated in Table S1) and cloned into the pXYFPN4 vector.

#### Strains constructions

Three categories of strains were constructed: (1) gene deletions or gene replacements were constructed using pNPTS138-derived plasmids (listed in Table S2) via two homologous recombination events: first, the suicide vectors were integrated at the desired location into the genome via homologous recombination and selection for kanamycin resistance. Excision of the suicide vector via a second homologous recombination event was then selected by plating on sucrose (0.3%)-containing PYEA plates and the isolated colonies were subsequently selected for kanamycin sensitivity. The mutants were identified by PCR using primer pairs used to construct pNPTS138 derivatives (Table S1) or by DNA sequencing for gene replacements; (2) pXYFPN4 derivatives were integrated at the native *xylX* locus by transformation. Positive clones were screened using RecUni-1 and RecXyl-2 primers (1). These strains were used for the inducible expression of MutS, MutL, UvrD derivatives in the presence of 0.3% xylose; (3) In JC1724, JC1845 and JC2211, the pNPTS138-DnaQ-G13E plasmid was integrated at the native *dnaQ* locus. The resulting strains express the *dnaQ(G13E)* allele together with a truncated and inactive *dnaQ*' allele. Oligos TC34 and M13-F were used to check the integration sites of the plasmid.

#### Immunoblot analysis

YFP-tagged proteins were resolved on 8% SDS-PAGE gels. Proteins were then transferred to PVDF membranes (Millipore). Two antibodies were used for immunodetection: Anti-GFP antibody (GFP Tag Antibody, GF28R, Mouse IgG, Thermo Fisher) was diluted 1:3000 and Anti-Mouse IgG HRP Conjugate (Promega) was diluted 1:5000. The signal was visualized using a chemiluminescent reagent (Amersham ECL Prime Western Blotting Detection Reagent, GE) and a FUSION FX (VILBER) scanner. Images were processed by Photoshop.
